# The Extreme Diversity Of Retinal Amacrine Cells Has Deep Evolutionary Roots

**DOI:** 10.64898/2026.03.07.710289

**Authors:** Dario Tommasini, Aboozar Monavarfeshani, Vishruth Dinesh, Joshua Hahn, Jared Tangeman, Olivier Marre, Seth Blackshaw, Teresa Puthussery, Joshua R. Sanes, Karthik Shekhar

**Affiliations:** Department of Neuroscience, University of California, Berkeley, CA, USA; Helen Wills Neuroscience Institute, University of California, Berkeley, CA, USA; Department of Molecular and Cellular Biology and Center for Brain Science, Harvard University, Cambridge, MA, USA; Department of Chemical and Biomolecular Engineering, University of California, Berkeley, Berkeley, CA, USA; Solomon H. Snyder Department of Neuroscience, Johns Hopkins University, Baltimore, Maryland, USA; Institut de la Vision, Sorbonne Université, INSERM, CNRS, Paris, France; Herbert Wertheim School of Optometry and Vision Science, University of California, Berkeley, CA, USA; Vision Sciences Graduate Program, University of California, Berkeley, CA, USA; Center for Computational Biology, University of California, Berkeley, CA, USA; Biophysics Graduate Group, University of California, Berkeley, CA, USA; Biological Systems Division, Lawrence Berkeley National Laboratory, Berkeley, CA, USA

## Abstract

Amacrine cells (ACs) comprise a heterogeneous class of inhibitory neurons in the vertebrate retina, exhibiting morphological and functional complexity rivaling that of cortical interneurons. Here, we integrate single-cell and single-nucleus transcriptomic atlases from 24 vertebrate species to reconstruct the evolutionary origins of this extreme diversity. We identify 42 orthologous AC types (oACs), most of which exhibit a one-to-one correspondence across amniotes and, in many cases, across vertebrates. While core molecular identities are conserved, AC types vary in abundance and gene expression across species, likely reflecting adaptations to distinct visual ecologies. AC diversity scales with that of retinal ganglion cells (RGCs), indicative of co-evolution. Finally, we suggest that ACs arose from an AC-RGC hybrid precursor, with glycinergic ACs diverging early in vertebrate evolution, followed by a bifurcation between RGCs and GABAergic ACs. Together, these findings establish a unified evolutionary framework for understanding the diversity, development, and function of a class of inhibitory neurons across vertebrates.

## Introduction

The vertebrate retina is a layered neural circuit that initiates visual processing by transforming light into patterned electrical signals for interpretation by the rest of the brain (*1*). Its basic blueprint, conserved across vertebrates for over 500 million years (*2–5*), consists of five major neuronal classes: photoreceptors (PRs), bipolar cells (BCs), ganglion cells (RGCs), horizontal cells (HCs), and amacrine cells (ACs) (*1*, *6*). PRs detect photons and relay signals to BCs, which in turn drive RGCs, the retina’s sole output neurons. This feedforward pathway is shaped by two classes of inhibitory interneurons – HCs in the outer retina and ACs in the inner retina – that modulate the spatial and temporal aspects of RGC responses. Somata of ACs, on which we focus here, reside in either the inner nuclear layer (INL) or ganglion cell layer (GCL). Their dendrites stratify within the inner plexiform layer (IPL), where they synapse with BCs, RGCs, and other ACs. Each of the five neuronal classes comprises multiple types distinguished by their morphology, physiology, connectivity, and molecular profile (*7*). Connectivity between and within classes is highly specific, and this specificity makes each RGC type selective for a small subset of visual features, such as edges, motion, or color contrasts, which they transmit to central brain regions (*1*).

Among the five classes of retinal neurons, ACs are the most heterogeneous (*8–10*). Their striking morphological diversity was initially revealed by histological and immunohistochemical analyses (*8*, *10–12*). This view has been reinforced by modern approaches, including connectomic reconstructions from electron microscopy (*13*), single-cell transcriptomics (*14–16*), and large-scale functional imaging (*17*), all of which underscore their extensive heterogeneity (**Fig. 1A**, **Table S1**). AC diversity plays critical roles in shaping feature selectivity in visual computations, owing to the specific connectivity of each type with a subset of BC and RGC types (*18*, *19*).

**Fig. 1.**
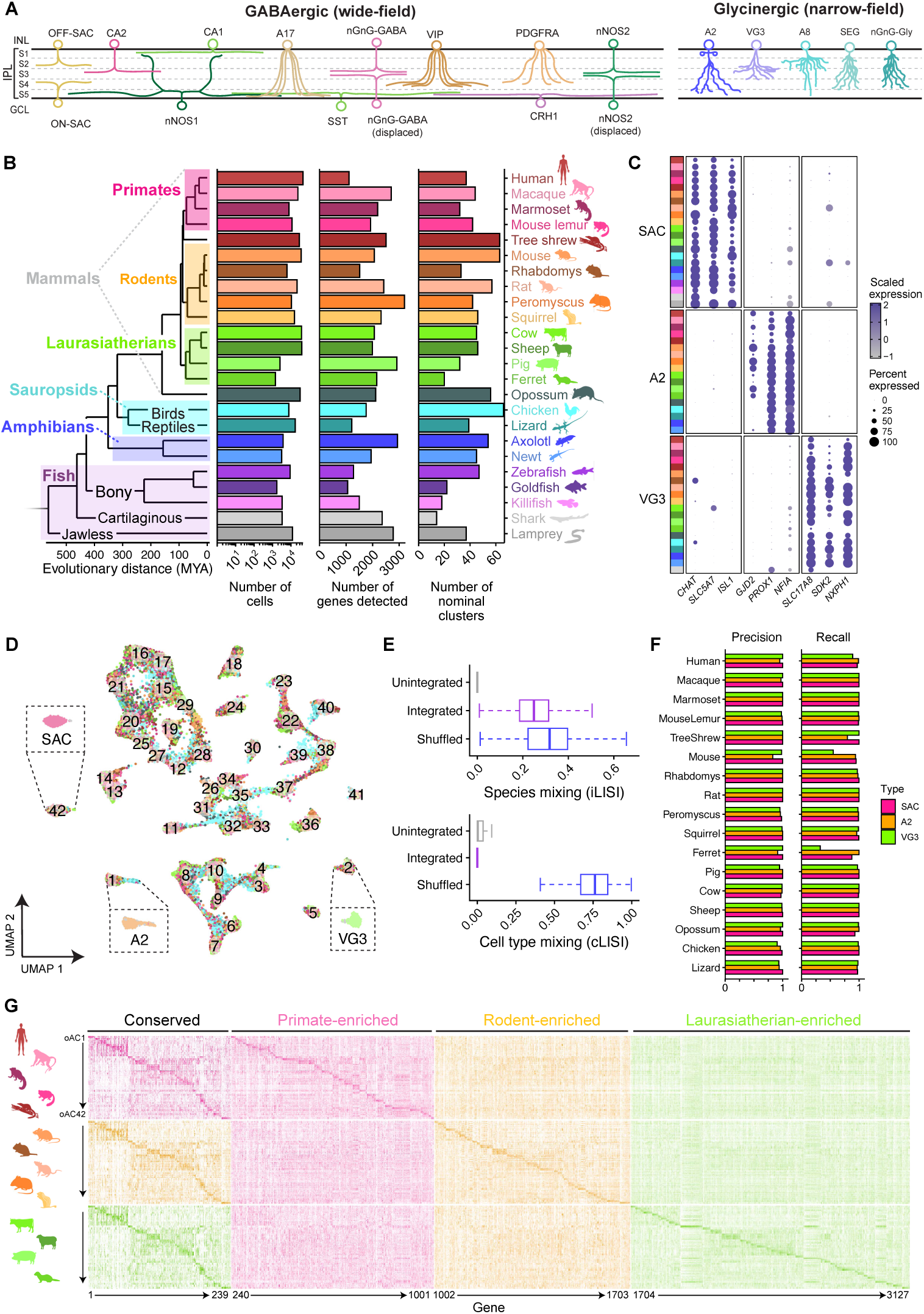
Amniote amacrine cell orthotypes (oACs). A) Morphological diversity of some characterized AC types. Sketches are based on references in **Table S1**. B) Phylogeny of species studied. Bar graphs on the right show number of cells (log scale), average number of genes detected, and the number of nominal clusters. Major phylogenetic clades are highlighted with boxes. C) Dotplots showing expression patterns of SAC, A2, and VG3 AC markers identified by independent manual annotation within each atlas. Species indicated by colors used in **B**. D) Uniform manifold approximate projection (UMAP) embedding of integrated amniote ACs, with cells mixing across species. Cells are colored by species of origin, following the color scheme from Fig. 1B. Insets highlight clusters corresponding to manually annotated SAC, A2, and VG3 ACs from Fig. 1C. oACs are numbered 1 through 42. E) Integration metrics. Integration local inverse Simpson index (iLISI) metric shows high level of species mixing in the integrated space. Celltype LISI (cLISI) shows minimal mixing of distinct cell types (SAC, A2, VG3) in the integrated space. The shuffled values were calculated from an integration of shuffled expression data. F) Precision and recall values for the three manually annotated types in each amniote species. G) Heatmaps showing conserved and clade-enriched genes for primates (top rows), rodents (middle rows), and laurasiatherians (bottom rows). The y-axis represents cells ordered by orthotype within each clade. The x-axis represents conserved, primate, rodent, and laurasiatherian-enriched genes, all of which are listed in **Table S3**.

The extraordinary complexity of ACs presents formidable challenges for classification. In the mouse, for example, single-cell transcriptomics has identified over 60 transcriptionally distinct AC types, representing nearly half of all neuronal diversity in the retina (*14*, *20*). Yet, fewer than 20% of these types have been characterized in more than one modality, and fewer still have been traced across species (*7*). As a result, fundamental questions about the evolution of AC types—their conservation, diversification, and origin—remain unresolved.

Recent advances in single-cell transcriptomics offer a new way forward. Just as genome evolution can be studied by aligning DNA sequences, cell type evolution can be inferred by aligning gene expression profiles across species. We and others have shown that such approaches can reveal conserved neuronal types across large evolutionary distances (*15*, *21–23*). In prior work, we used single-cell and single-nucleus RNA-sequencing (sc/snRNA-seq) to map the conservation of PRs, HCs, RGCs and BCs across 17 vertebrate species, identifying many orthologous types conserved across tetrapods (*21*). Targeted studies in lamprey and zebrafish support the ancient origins of some cell types and circuits (*24*, *25*). However, a comprehensive evolutionary analysis of ACs has been lacking.

To address this gap, we have reconstructed the molecular evolution of AC diversity using sc/snRNA-seq data from 24 vertebrate species. Integrative genomic analysis revealed 42 orthologous AC types (oACs), most of which are conserved across tetrapods and many of which are traceable to the vertebrate common ancestor.

This unified atlas both refines the classification of well-studied AC types and enables cross-species annotation for many previously uncharacterized types, grounding their identities in a shared evolutionary origin. We further identify a conserved transcription factor (TF) code that may govern the diversification of AC types. In addition, we show that AC diversity scales with that of RGCs, pointing to concerted evolution of pre- and postsynaptic partners.

Finally, we present evidence for divergent evolutionary trajectories of the two major AC subclasses: glycinergic ACs, which are typically narrow-field and vertically integrate signals across IPL sublaminae, and GABAergic ACs, which are usually wide-field and mediate lateral interactions across the retina (*6*, *12*, *19*, *26*). Our comparative analysis indicates that glycinergic ACs diverged early in vertebrate history from an AC/RGC hybrid ancestor. Subsequently, GABAergic ACs diverged from RGCs. This scheme supports a shared ancestral origin for ACs and RGCs and provides an explanation for why several GABAergic AC types exhibit RGC-like features.

## Results

### Amacrine cell atlases across 24 vertebrates

We previously assembled sc/snRNA-seq atlases from 17 vertebrate species: sea lamprey (*25*), zebrafish (*24*, *27*), anole lizard (*21*), chicken (*28*), and 13 mammals (*21*). Here, we expanded this dataset for AC analysis by profiling four additional species—mouse lemur, rat, axolotl, and Iberian ribbed newt (**Fig. S1**)—and incorporating published data from three others—killifish (*29*), goldfish (*30*), and catshark (*31*)—for a total of 24 species (**Fig. 1B; Table S2**). As ACs typically represent <10% of retinal cells (*1*, *32*), we used a combination of antibody-based selection and depletion approaches to enrich for ACs in some collections (see **Methods**). For zebrafish, where our original dataset contained too few high-quality ACs, we incorporated transcriptomes of ∼11,500 ACs from Ref. (*33*).

For each species, we first identified ACs based on the expression of canonical markers (*PAX6*, *TFAP2A*/*B*/*C* and *GAD1/2* or *SLC6A9*), and then divided them into molecularly distinct clusters using dimensionality reduction and graph-based clustering, following established workflows (*14*, *34*). After filtering low-quality cells, doublets, and contaminating cell classes, the resulting datasets contained a total of 375,881 high-quality ACs, with 1,434–44,493 cells per species (**Fig.** Error! Reference source not found.**B**). The number of nominal clusters varied with cell count, ranging from 14 in cat shark (N=3,333 cells) to 66 in chicken (N=7,601 cells).

Inspection of marker gene patterns (**Table S3**) enabled us to relate several of these clusters to well-studied AC types from rodents and primates, such as starburst (SACs), A17, catecholaminergic (CA1/2), A2, <u>S</u>atb2^+^<u>E</u>bf3^+^<u>G</u>lyT1^+^ (SEG), and VGLUT3^+^ (VG3) ACs (*6*, *14*, *18*). In particular, SAC, A2, and VG3 ACs, which have multiple reliable molecular markers, could be annotated confidently and were detected in nearly every species (**Fig.** Error! Reference source not found.**C**).

### Integration across amniotes identifies 42 amacrine cell orthotypes (oACs)

In contrast to these well-known types, cross-species homology was not apparent for most AC clusters. This motivated a data-driven integration to achieve consistent and evolutionarily grounded AC annotations. To this end, we first focused on the 17 amniotes, excluding the amphibians and fish due to their greater evolutionary distance. We integrated the transcriptomes based on the shared expression patterns of 6,305 one-to-one gene orthologs (*21*, *35*). Before integration, cells primarily clustered by species of origin (**Fig. S2A**), but after integration, cells were well-mixed across species (**Fig. 1D**, left panel). The integration was performed without reference to intra-species cell type labels, yet known types such as SAC, A2, and VG3 co-clustered post-integration (**Fig. 1D**, insets), suggesting accurate alignment. The integrated embedding retained internal structure, consistent with AC heterogeneity, but this structure disappeared when expression values were randomly shuffled (see **Methods**), ruling out integration artifacts (**Fig. S2A).** We quantified integration quality by calculating the local inverse Simpson index (LISI) (*36*), a statistical measure of mixing in the embedded space, at the level of species labels (iLISI) as well as cell type labels (cLISI). The values were indicative of high inter-species mixing and low inter-cell type mixing (**Fig. 1E**).

Unsupervised clustering of cells in the integrated space yielded 42 clusters defined by shared molecular signatures (**Fig. 1D)**, all of which expressed canonical AC markers and included cells from multiple species (**Fig. S2B)**. Following conventions from our earlier study (*21*), we term these integrated clusters AC orthotypes (oAC1-42), each corresponding to a conserved, molecularly defined type. To assess the fidelity of the integration, we tested how well the integrated clusters recovered the manually annotated SAC, A2, VG3 cells shown in **Fig. 1C**. The correspondence was high, with 98% precision (95% CI: 0.97–0.99) and 96% recall (95% CI: 0.93–0.99) (**Fig. 1F**). The rare instances of lower recall were attributable to underclustering in the species-specific atlas rather than integration artifacts. For example, ferret SACs failed to form a discrete cluster in the ferret atlas due to low sampling, but were recovered in the SAC oAC as six *SLC5A7+* cells, likely representing bonafide SACs (*14*) (**Fig. S2C**). Importantly, oAC definitions were robust across a range of integration parameters (**Fig. S2D)** and were reproducible using a different integration method, Harmony (*36*) (**Fig. S2E**). oACs also aligned well with AC types from the mouse and primate atlases (*14*, *15*) (**Fig. S3A**), which have undergone extensive experimental validation.

We next inspected differentially expressed genes (DEGs) associated with each oAC for evidence of conserved transcriptional signatures (see **Methods**), as would be expected for true orthotypes. We found DE markers for each oAC within primates, rodents, and laurasiatherians, thereby controlling for clade-specific expression differences. A comparison of the three clade-specific DEG sets revealed a core set of type-specific markers conserved across all three clades, as well as genes selectively enriched in each of the three orders or superorders (**Fig. 1G**, **Table S3**). Together, these results demonstrate that ACs can be accurately aligned across species based on conserved transcriptional programs, providing a robust foundation for further analyses.

### Neurotransmitter identity of oACs

Most AC types use either GABA or glycine as their primary neurotransmitter, often along with amines and neuropeptides (*7*, *8*). We assessed the expression of GABAergic (*GAD1, GAD2, SLC6A1, SLC6A13, SLC6A11*) and glycinergic markers (e.g., *SLC6A9, SLC6A5*) across oACs. In samples collected without antibody-based selection of ACs, glycinergic and GABAergic cells were present in roughly equal numbers, consistent with their native proportions (*37–39*). However, there were more GABAergic types than glycinergic types within each species, echoing previous morphological and molecular studies in rodents (*8*, *14*).

Expression of *GAD1/2* (GABA synthesis) and *SLC6A9* (glycine transporter 1) was mutually exclusive across oACs, yielding 10 glycinergic and 32 GABAergic oACs (**Fig. 2A).** Thus, all oACs fell into either the glycinergic (oAC1-10) or GABAergic (oAC11-42) subclass. Notably, the integration also revealed clear oAC assignments for the mouse types initially termed non-GABAergic, non-glycinergic (nGnG) based on their low expression of both GABAergic and glycinergic markers (*40*). The four mouse nGnG types (*14*) mapped to two oACs, GABAergic oAC36 and glycinergic oAC8 (**Fig. 2B; Fig. S3A**). Neurotransmitter gene expression patterns in these types were species-dependent: low in mouse, peromyscus, and rat (consistent with their initial assignment as nGnG based on immunohistochemistry), but elevated in several other species (**Fig. 2C**). Similarly, two additional orthotypes, oAC9 and oAC10, showed potential dual GABA/glycine signatures in certain species. These findings suggest that nGnG and dual-transmitter identities may represent species-specific specializations, whereas at the orthotype level, oACs are either GABAergic or glycinergic.

**Fig. 2.**
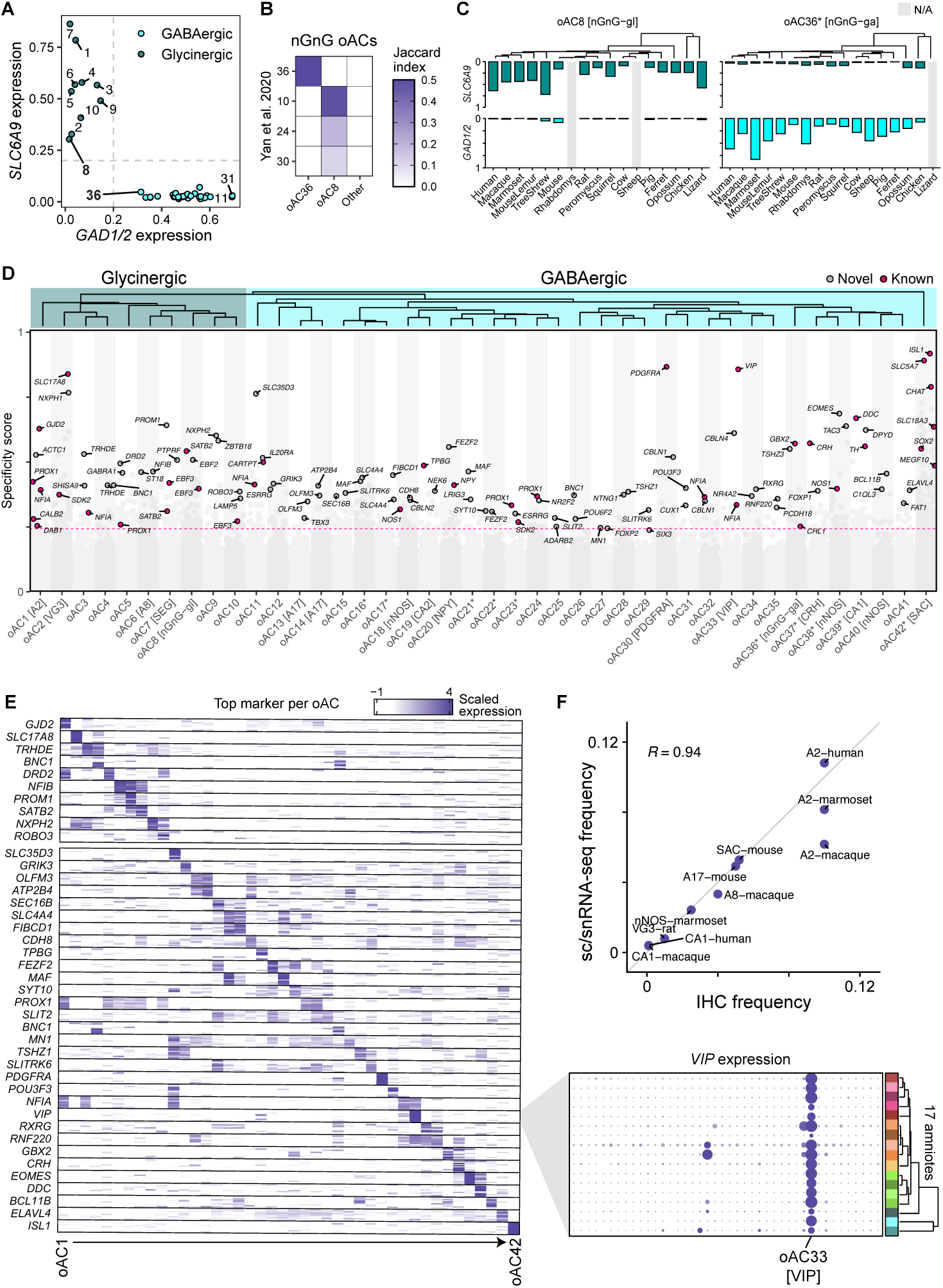
Classification and annotation of amniote oACs. A) Scatterplot showing *GAD1/2* expression and *SLC6A9* expression (as defined in **Methods**) for 42 oACs, which separate into 32 GABA and 10 glycinergic types. B) 4 nGnG ACs from mouse (*14*) map to 2 oACs with the lowest *GAD1/2* and *SLC6A9* expression in their subclass (GABAergic oAC36 and glycinergic oAC8). C) GABAergic and glycinergic scores for nGnG oACs across species reveal that low *GAD1/2* and *SLC6A9* expression is variable across species and particularly low in mouse. oAC8 [nGnG-gl] is glycinergic in most species, whereas the oAC36* [nGnG-ga] is GABAergic in most species. Grey bars indicate species in which these oACs were not confidently identified. D) Specificity score (y-axis, see **Supplementary Materials and Methods**) for key genes in each oAC (x-axis). Each dot represents a gene-oAC combination. The top two markers for each oAC are labeled, as well as known markers that passed the significance threshold (p < 2.5×10^-3^ by permutation test, 718,746 permutations). Other markers are represented as gray dots. oACs were annotated based on known markers. The dendrogram is a hierarchical clustering tree (with complete linkage) using Pearson correlation distance (1 - R) between oACs and 2000 highly variable genes. E) Heatmap showing the expression patterns of top oAC markers from Fig. 2D. Each rows in the heatmap represents the expression of a gene in a given amniote, ordered as shown in the phylogenetic tree in the inset. Each column represents an oAC, ordered as in Fig. 2D. Inset shows the expression of *VIP* in oAC33 [VIP]. F) Scatterplot showing tight correspondence (pearson R = 0.94) between relative frequencies of AC types based on sc/snRNA-seq (y-axis) and IHC (x-axis). References for IHC quantification are listed in **Table S1**.

### Evolutionarily grounded annotation of oACs

We next annotated oACs based on conserved molecular markers, drawing on prior histological and molecular data from well-studied species such as mouse, rat, rabbit, macaque, and human (*7*, *18*). For each oAC, we prioritized genes that were both highly conserved across species and strongly selective in their expression patterns. The top-ranked markers (**Table S3)** based on specificity scores for each oAC are shown in **Fig. 2D**, and the expression patterns of a subset are displayed in **Fig. 2E**.

Several oACs mapped to well-studied AC types (**Table 1**; **Fig 2D**). These included SACs (oAC42) marked by *SLC5A7*, *CHAT*, and *ISL1*; A2 ACs (oAC1) expressing *GJD2* and *PROX1*; SEG ACs (oAC7) marked by *SATB2* and *EBF3*; catecholaminergic CA1 ACs (oAC39) expressing *DDC* and *TH*; CA2 ACs (oAC19) expressing *TH* and *TPBG*; VG3 (oAC2) marked by *SLC17A8*; A17 ACs (oAC13 and oAC14), co-expressing *PRKCA* and *SDK1* (**Fig. S3B,C)**; the *CRH*+ GABAergic AC (oAC37) (*41*); the *GBX2*+ nGnG AC (oAC36) (*14*, *41*); and A8 ACs (oAC6), based on robust *SYT2* expression in macaque (*42*), which was conserved in rat, squirrel, and opossum but not in other species.

**Table 1.**
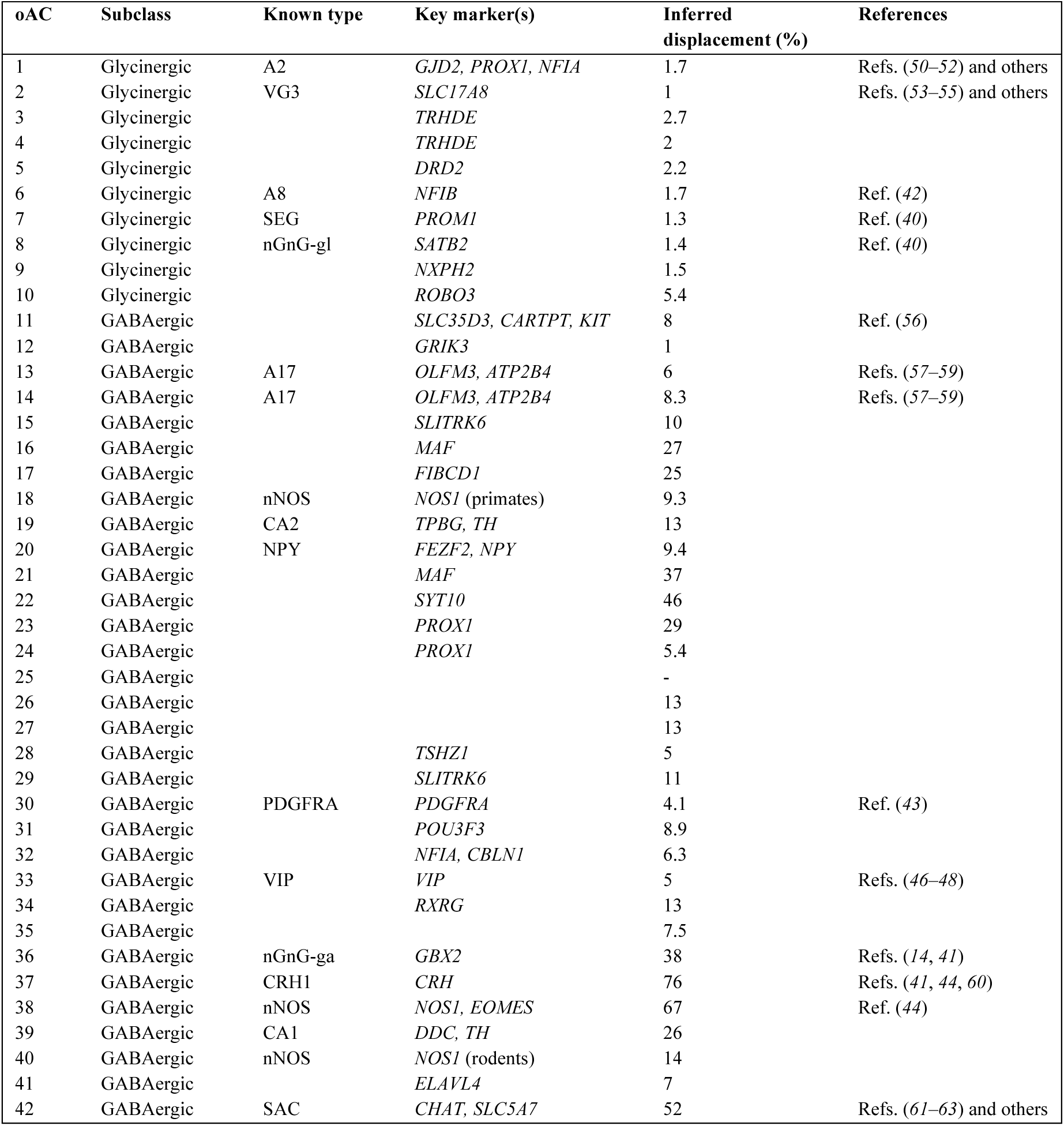
Table of amniote oACs showing subclass, putative type or highly conserved marker, and inferred displacement. Displacement was inferred from a cross-sectional mouse spatial transcriptomic atlas. (***64***)**; oACs with greater than 20% are indicated with an asterisk. We did not infer displacement for oAC25 because too few cells from this oAC were mapped to mouse.**

The integration also facilitated annotation of several conserved but poorly characterized types. For instance, oAC30—labeled by *PDGFRA* (**Fig. 2E**)—likely corresponds to a wide-field GABAergic type described previously (*43*), while oAC20, marked by *NPY*, likely corresponds to the NPY-releasing AC (*7*). Notably, the integration revealed three distinct neuronal nitric oxide synthase (nNOS) expressing AC types: oAC38, which expressed *NOS1* across mammals; oAC18, which expressed *NOS1* primarily in primates; and oAC40, which expressed *NOS1* primarily in rodents. oAC38 may correspond to the NOS-1 type reported previously (*44*), and was also marked by *EOMES* (**Fig. 2E**). In contrast, *NOS2* was not reliably detected in any species.

Integration also helped resolve certain ambiguous annotations in the species-specific atlases. For example, although *CARTPT* had been previously detected in macaque A17 ACs (*45*), this expression was restricted to macaque. Instead, *CARTPT* was an evolutionarily conserved marker of oAC11, whose top-ranked marker was the solute carrier *SLC35D3* (**Fig.** Error! Reference source not found.**D**). In addition, *VIP* expression was highly conserved in oAC33, but its expression in oAC19 and oAC32 was species-specific, consistent with morphological evidence for multiple *VIP+* types in rodents (*44*, *46–48*). Finally, while *TH* (tyrosine hydroxylase) is a rodent-specific marker of CA2/TH2 ACs (oAC19), it was also expressed in primate oAC33 (*VIP*+), highlighting species-specific differences that are not apparent when analyzing individual atlases in isolation.

While most AC somas reside in the INL, the somas of displaced ACs – a subset of GABAergic ACs – are located in the GCL. To identify which orthotypes may contain displaced ACs, we inferred soma displacement by transferring labels from a recent spatial transcriptomic atlas of the mouse retina (*49*), which identified 13 mouse types with greater than 20% displacement (**Table 1**). These mouse types converged onto 10 oACs, all of which were GABAergic (**Fig. S2D**)—oAC16, 17, 21, 22, 23, 36, 37, 38, 39, and 42. An additional 7 types, all GABAergic, exhibited >10% displacement.

In total, 9/32 GABAergic oACs and 5/10 glycinergic oACs could be mapped to securely known types based on expression patterns of known markers (**Table 1**). Throughout the remaining text, we refer to oACs by their number along with their type identity when assigned to a well-defined type (e.g., oAC42* [SAC]) or a defining evolutionarily conserved marker (e.g., oAC30 [PDGFRA]). oACs that are >20% displaced are indicated by an asterisk (*).

### Cross-species histological validation of select AC orthotypes

To assess the validity of the transcriptomic assignments, we used two immunohistochemical methods. First, we compared the cell type frequencies estimated by our atlases to cell counts from previous histological surveys (**Table S1**). As shown in **Fig. 2F**, there was strong agreement between transcriptomics and IHC (Pearson R = 0.94). Second, we immunolabeled novel markers (**Fig. 2D**) to identify four oACs in various mammalian retinas. oAC2 [VG3], which expresses the atypical vesicular glutamate transporter VGluT3, was positive for the novel marker *NXPH1*, which encodes a secreted glycoprotein involved in synaptic stabilization (**Fig. 3A**). The transcription factor *MAF* is expressed in two GABAergic oACs (oAC16* and oAC21*), both predicted to contain displaced cells. In agreement with this, MAF^+^ cells were observed in both the INL and GCL in mouse and macaque retinas. Approximately 30% of MAF^+^ cells resided in the GCL—consistent with proportions observed in spatial transcriptomics (**Fig. 3B**; **Table 1**). We also co-labeled EOMES and nNOS to identify oAC38* [nNOS], which is a wide-field, polyaxonal type that resembles the previously described nNOS-1 population (*44*) (**Fig. 3C**). EOMES^+^nNOS^+^ cells were mostly found in the GCL. To assess the conservation of oAC30 [PDGFRA], we stained retinas from mouse, guinea pig, rabbit, and human with antibodies against PDGFRA. In all four species, PDGFRA selectively labeled an INL-restricted GABAergic AC population (**Fig. 3D**), consistent with our prediction (**Table 1**). Notably, oAC30 is enriched in higher primates (**Fig. 3E**), a trend not seen in any other oAC (**Fig. S4, S5**).

**Fig. 3.**
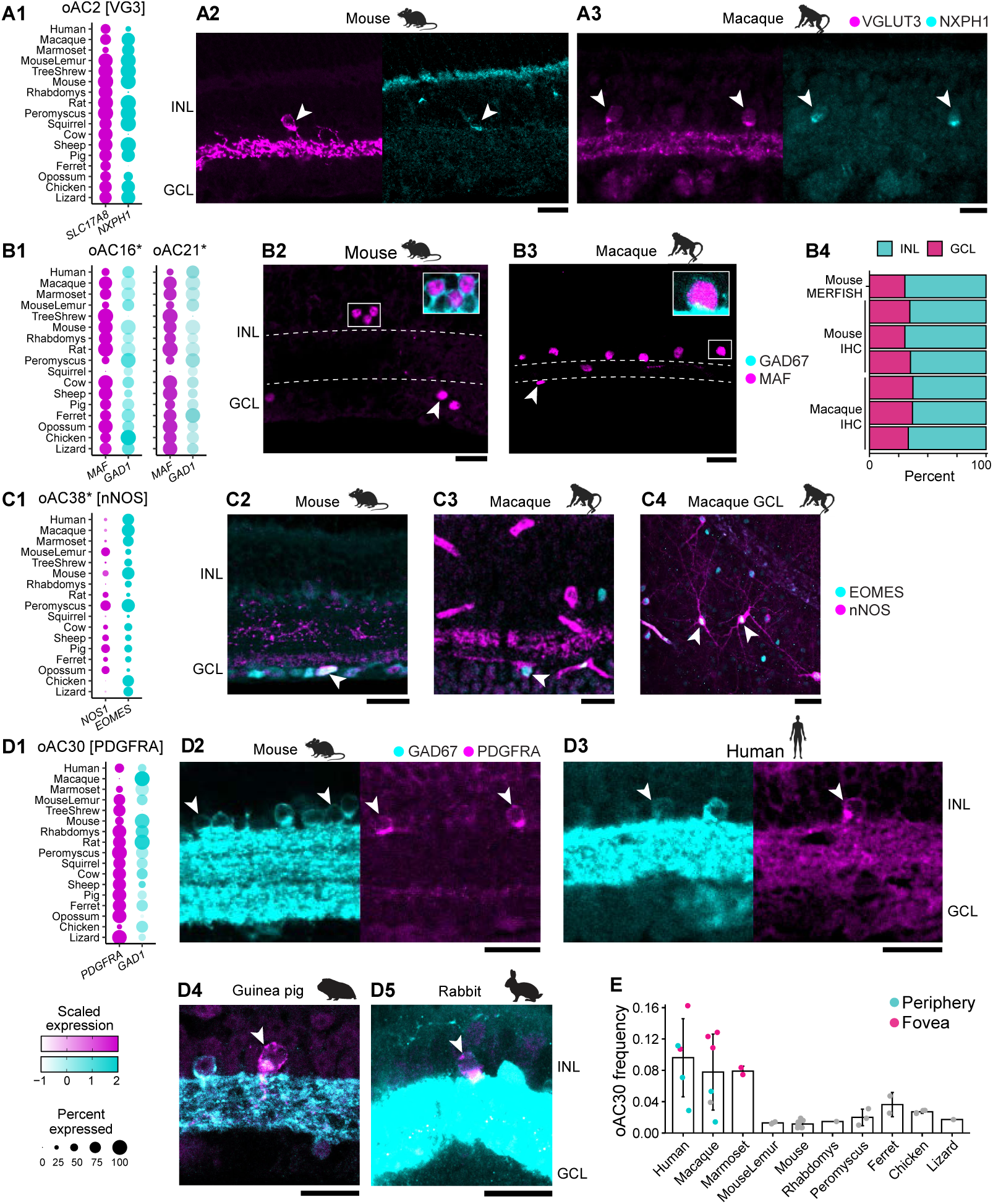
Histological validation of oACs. A) oAC2 [VG3]. A1) Dotplot showing coexpression of *SLC17A8* (gene encoding VGluT3) and *NXPH1* expression in oAC2 [VG3]. A2) Co-staining of VGluT3 and NXPH1 in a mouse retinal section. A3) Co-staining of VGluT3 and NXPH1 in a macaque retinal section. Double positive cells are shown by arrowheads. B) oAC16* and 21*. B1) Dotplot showing coexpression of *MAF* and *GAD1* in GABAergic oAC16* and oAC21*. B2-B3) Co-staining in mouse and macaque sections showing that MAF labels displaced GAD67+ GABAergic ACs. B4) Comparison of displacement of MAF+ somata between IHC and MERFISH. Choi et al.: 30% displaced; IHC in mouse: 33% based; IHC in macaque: 35%. n = 3 biological replicates per species; 196 cells counted total. Arrowheads point to examples of displaced MAF+ ACs. C) oAC38* [nNOS]. C1) Dotplot showing coexpression of *EOMES* and *NOS1* in oAC38* [nNOS]. C2-C4) EOMES and nNOS staining in mouse and macaque labels displaced, wide-field ACs (arrowheads). For C4, scale bar shows 50 µm. D) oAC30 [PDGFRA]. D1) Dotplot showing coexpression of *PDGFRA* and *GAD1* in oAC30 [PDGFRA]. D2-D5) PDGFRA and GAD staining in mouse, rabbit, guinea pig, and human. Arrowheads show double positive cells. E) Relative abundance of oAC30 [PDGFRA] in each transcriptomic atlas, measured as proportion of all ACs. Each dot represents a biological replicate. oAC30 showed the highest enrichment in primates out of all 42 oACs (**Fig. S4**, **S5**). In higher primates, pink dots represent foveal samples and cyan dots represent peripheral samples. Only samples without antibody-based selection were used for quantification. Dotplot legends are shown at the bottom left of the figure. All scale bars indicate 20 µm unless specified. Arrowheads mark the oAC of interest unless specified.

### Conservation of AC diversity across amniotes

We next evaluated how each of the 42 oACs corresponded to species-specific clusters (**Fig. S6, S7A,B**). A strong correspondence within every species would support the notion of a cell type inherited from the last common ancestor, whereas correspondence in only a subset of species could suggest lineage-specific gains or losses. We quantified this correspondence using a “presence score”, defined as the Jaccard index between an oAC and its best matching cluster within each species. A presence score of 1 indicates that the oAC is represented in the species by a single AC cluster, whereas values near zero indicate that no corresponding cluster is detected. Intermediate scores indicate partial correspondence, consistent with the presence of the oAC but without a 1:1 mapping to a single cluster—for example, if the cell type occurs at low frequency or is split across multiple subtypes.

We found that ∼80% of oACs (32/42) could be confidently identified in >80% (12/15) of the mammalian species (FDR < 0.10, see **Methods**) (**Fig. S7C)**. oACs with weaker support tended to be rare and sparsely sampled (**Fig. S7D**), suggesting that their apparent absence is more likely due to sampling than true evolutionary loss. In fact, even these weakly supported orthotypes could be reliably detected in 5 or more species (**Fig. S7D**). Taken together, these results suggest that AC types are orthologous across mammals (**Fig. 4A**), with occasional merging of related oACs and dropout of rare, lowly sampled types.

**Fig. 4.**
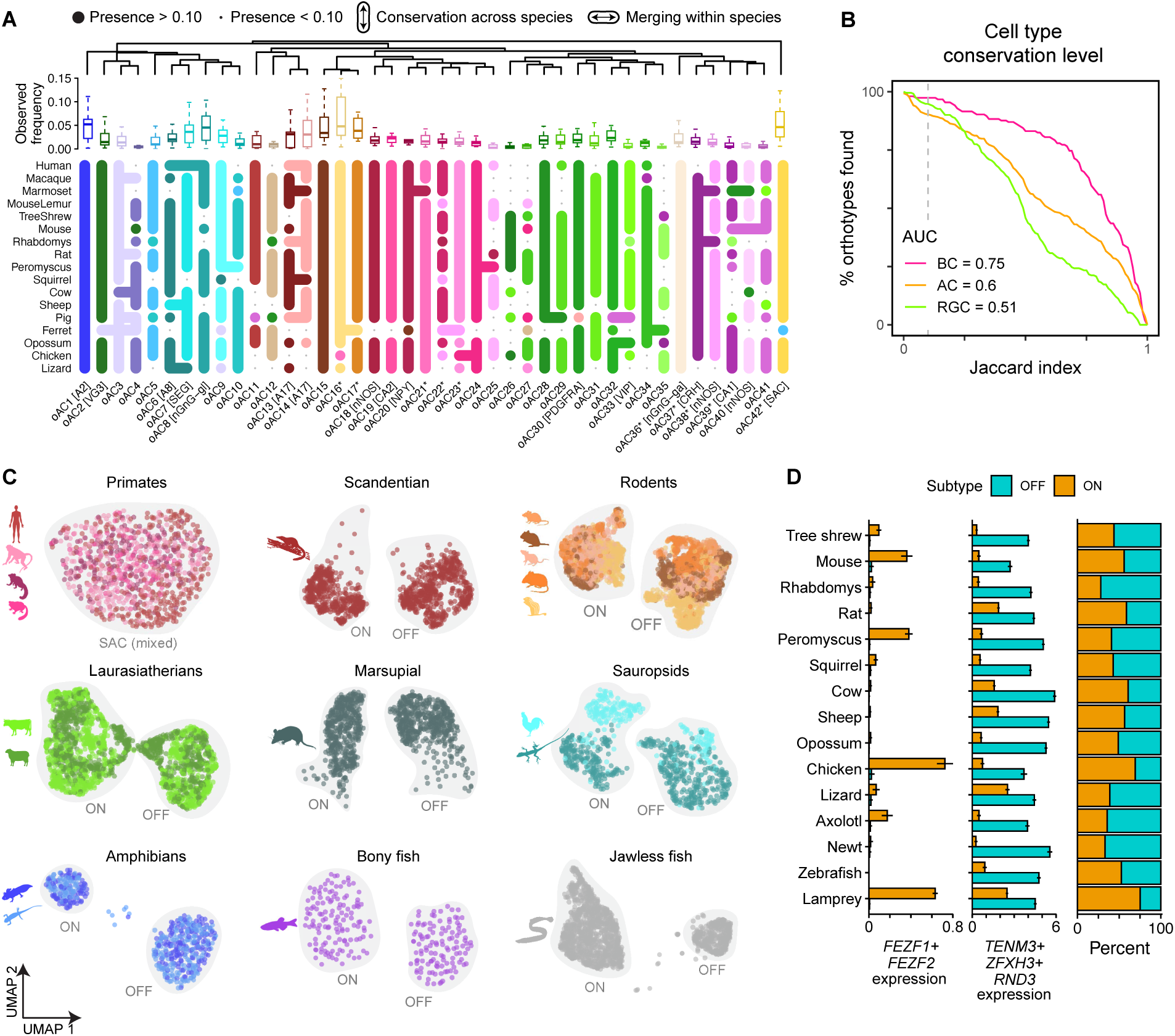
Conservation of oACs across amniotes. A) Snake plot showing the correspondence of oACs across amniotes. Colors of dots correspond to oACs (color key for oACs is the same as in **Fig. S2)**. Vertical connections indicate cross-species homology while horizontal connections show merging of related orthotypes within a species (see **Methods**). Boxplot above shows the observed (raw) frequency of each orthotype across 17 amniotes, revealing that orthotypes absent in some species tend to have low global frequency, consistent with their apparent absence arising from sampling limitations rather than true loss. See **Fig. S7C** for more information. B) Line graph showing the percentage of oACs, oBCs and oRGCs resolved at different Jaccard index thresholds. We computed the area under this curve to quantify cross-species conservation across a common set of 11 species shared across all three multi-species datasets to ensure comparability. Dashed vertical line indicates the threshold used for Fig. 4A. C) UMAPs showing subclustered ON and OFF SACs across primates, rodents, laurasiatherians, scandentian (tree shrew), marsupial (opossum), sauropsids, amphibians, a bony fish (zebrafish), and a jawless fish (lamprey) (see **Methods**). D) *Left and middle*: Box plots showing the expression of known markers of ON and OFF SACs used for annotation. *Right*: stacked bar graph showing the relative abundance of ON and OFF SACs in each species. Note that ON and OFF SACs could not be resolved in any of the primates (see top left panel in Fig. 4C).

To assess the extent of molecular conservation of AC types relative to BCs and RGCs, we calculated the percentage of oACs resolved at different Jaccard index thresholds and compared it to oRGCs and oBCs from our previous study (*21*). Overall, the extent of molecular conservation of AC types, while lower than that of the highly conserved BCs, is higher than that of RGCs (**Fig. 4B**), despite being much more diverse and therefore more susceptible to sampling artifacts.

We highlight an interesting case wherein variation within an orthotype is conserved: oAC42* [SAC]. SACs can be subdivided into functionally distinct ON and OFF subtypes, which reside in the GCL and INL, respectively (*6*, *7*, *65*). ON and OFF SACs reliably segregated in 9/15 mammals (**Fig. 4C)** based on gene expression: *FEZF1* and *FEZF2* in ON SACs and *TENM3*, *ZFHX3*, and *RND3* in OFF SACs (*25*, *49*) (**Fig. 4D**). This separation was not evident in the four primates—even with a supervised classification (see **Methods**)—and SACs were too sparse in pig and ferret to be subclustered. SACs could, however, be subclustered into ON and OFF subtypes in sauropsids, amphibians, zebrafish and lamprey (*25*) (**Fig. 4C**).

Consistent with previous reports (*49*), gene expression profiles of ON and OFF SACs were highly similar in all species (Pearson *R* = 0.94–0.99), highlighting their transcriptional proximity despite their functional and morphological differences. Proportions of ON and OFF SACs varied widely across the phylogeny (**Fig. 4D**), likely reflecting species-specific adaptations.

### A conserved regulatory logic underlies amacrine cell heterogeneity

Although oACs are defined by statistically conserved gene expression patterns, substantial interspecies variation in gene expression exists within each oAC. This raises the question of how molecular and functional diversity are generated while preserving stable AC identities. An attractive possibility is that AC types are organized by a conserved regulatory logic, in which diversity arises through a limited number of coordinated transcription factor (TF) switches acting on a shared regulatory scaffold. To test this idea, we adapted a maximum parsimony framework from phylogenetics (*66*, *67*), reasoning that conserved TF expression patterns across AC types should minimize the total number of regulatory changes required to explain their relationships.

Consistent with this idea, TFs constituted an especially large fraction (62%) of highly conserved genes (score > 0.7) across amniote oACs (*p* < 10^-18^, hypergeometric test; **Fig. S8**).

To ask whether a transcriptional code could account for some aspects of AC diversification, we focused on a set of 70 conserved TFs whose expression could be cleanly binarized across species (see **Methods**). This TF set was enriched for factors previously implicated in AC biology in the literature (38/70 genes, *p* <10^-17^; hypergeometric test; PubMed accessed 9/2025), indicating that it captures known regulators of AC identity. Using these TFs, we constructed 132 maximum parsimony trees and derived a consensus topology (**Fig. 5A**, top). The resulting tree was highly robust to both TF resampling and alternative reconstruction methods (UPGMA, neighbor-joining, and maximum-likelihood). We hypothesize that clades defined solely by conserved TF expression correspond to developmentally related groups of oACs.

**Fig. 5.**
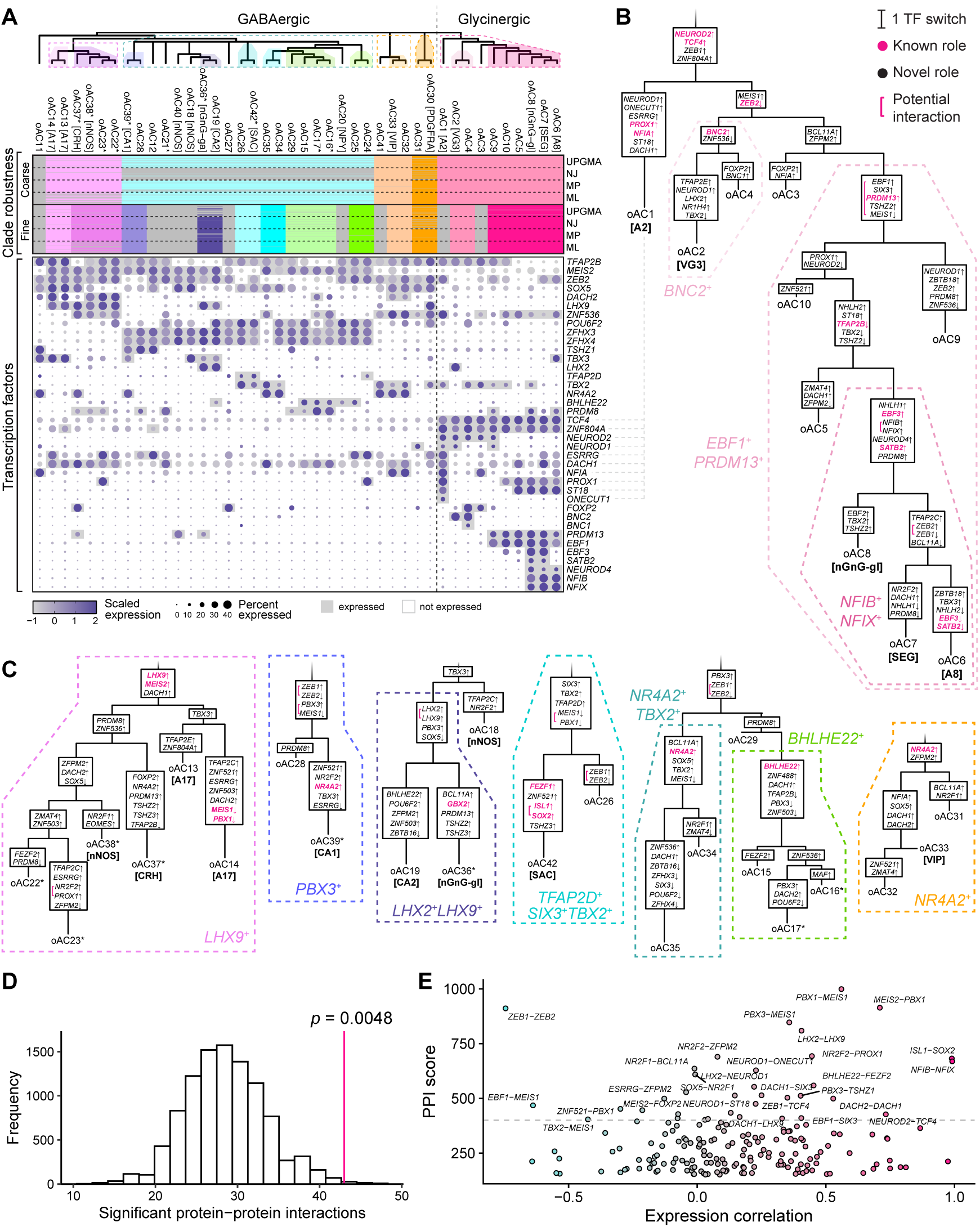
Conserved transcription factor codes in oACs. A) *Top*: Consensus maximum parsimony tree constructed based on expression patterns of 70 transcription factors (TFs) conserved across amniotes. The tree was partitioned into coarse (dashed boxes) and fine clades (filled boxes). *Middle*: Heatmaps showing robustness analysis of the trees to four alternative methods (UPGMA, neighbor joining [NJ], maximum parsimony [MP], and maximum likelihood [ML]) and removal of each TF (70 leave-one-out trees per method). Each row represents one cross-validation. Cells are colored if the corresponding clade was identified in that permuted tree. Upper and lower heatmaps correspond to the coarse- and fine-resolution clades, respectively. *Bottom*: Dot plots of conserved TFs that are expressed in a clade-selective fashion. After binarization, TFs were considered either expressed (gray box) or not expressed (white box) in each oAC. B) Reconstruction of most parsimonious sequence of TF switching events in the glycinergic clade (right dashed box in A). TFs whose role has been previously reported in AC development are highlighted in pink. C) Select clades among GABAergic oACs. Full parsimony tree is shown in **Fig. S9**. D) Histogram showing the null distribution of protein-protein interactions (PPI) when TF switching events are shuffled across the tree. The observed number of interactions (n = 43) is significantly enriched for PPI. E) PPI score from STRING database versus gene-gene expression correlation (computed using average log-transformed, normalized expression of oACs). Putative cooperativity events are positively correlated (e.g. *PBX1*/*MEIS1*), whereas putative antagonisms are negatively correlated (e.g. *ZEB1*/*ZEB2*). Dots are colored by expression correlation, with pink indicating positive correlation, and cyan indicating negative correlation.

We next reconstructed TF gain and loss events along the consensus tree (**Fig. S9**). Within the glycinergic clade, all types shared a core signature of two TFs, *TCF4* (*14*) and *ZNF804A*, upon which type-specific refinements were superimposed (**Fig. 5B**). The outgroup oAC1 [A2] uniquely co-expressed *NFIA* (*68*), *PROX1* (*69*), *ST18*, and *ONECUT1*. Next, differential activation of *MEIS1*, *BNC2* (*70*), and *BNC1* distinguished oAC3, oAC2 [VG3], and oAC4 from one another. The remaining six glycinergic types expressed *PRDM13*, *SIX3*, and *EBF1*. This subclade was further resolved by stepwise changes: 1) upregulation of *PROX1* accompanied by downregulation of *NEUROD2* (5 oACs), 2) upregulation of *ST18* with concomitant downregulation of *TFAP2B* (4 oACs), and 3) upregulation of *EBF3*, *SATB2,* and *NFIX/NFIB* (3 oACs). Notably, oAC6 [A8] was placed immediately adjacent to oAC7 [SEG], suggesting that the A8 type is closely related to SEG ACs but differs by loss of *SATB2*/*EBF3* and gain of *TBX3*.

We also identified several robust clades within the GABAergic ACs (**Fig. 5C, Fig. S9**), including an *LHX9*^+^ clade (6 types) (*71*, *72*), a large *ZFHX3/4*^+^ clade (20 types), and an *NR4A2*^+^ clade (3 types). Interestingly, the four oACs in the *LHX9*^+^ clade –oACs 13, 14, 37* [CRH], and 38* [nNOS] – all stratify in S5 (**Fig. 5C**, left), and three oACs in a *TBX3*^+^ clade – oAC18 [nNOS], oAC19 [CA2] and oAC36* [nGnG-ga] – are all predicted to stratify in S3 (*71*). This result raises the possibility that transcriptional programs regulate not only cell type identity but also laminar targeting of AC dendrites.

Finally, we find that putative co-regulated genes—defined as TFs inferred to change at the same point along the tree—were significantly enriched for putative protein-protein interactions from the STRING database (*73*) (*p* < 0.005 by permutation test, 10,000 permutations; **Fig. 5D**). This enrichment supports the interpretation that these coordinated switches represent functional regulatory modules. For instance, we predicted multiple instances of co-activation of *PBX1* with either *MEIS1* or *MEIS2* (**Fig. 5E**), consistent with their known cooperative DNA-binding interactions (*74*). We also detected multiple instances of mutual exclusivity between *ZEB1* and *ZEB2*, which are known to function antagonistically (*75*). Finally, co-activation of *NFIB* and *NFIX* was inferred in oAC6 [A8], oAC7 [SEG], and oAC8 [nGnG-gl], consistent with evidence that NFIB can activate *NFIX* expression *in vitro* (*76*).

### Conservation of AC diversity in basal vertebrates

To investigate whether AC types are conserved beyond amniotes, we extended our orthotype analysis to include two amphibians (axolotl and Iberian ribbed newt), three teleost bony fish (zebrafish, goldfish, and killifish), one cartilaginous fish (cat shark), and one jawless fish (lamprey). To bridge these analyses with the amniotes, we co-analyzed them with the two sauropsids (chicken, lizard), for a total of nine non-mammalian vertebrates. Because teleost fish underwent an additional whole-genome duplication event, we employed SAMap (*77*), a recently developed integration tool that incorporates complex gene homology relationships. As an initial validation, we applied SAMap to integrate the six major retinal cell classes (PR, BC, HC, AC, RGC, MG) across these nine species. We observed high conservation of cell classes, with class-level orthology supported by hundreds of homologous genes (**Fig. S10A, B**). At the cell type level, BCs, known to be highly conserved among mammals (*21*), also showed strong cross-species alignment among these non-mammalian species (**Fig. S10C,D**).

Application of SAMap to non-mammalian ACs revealed good interspecies mixing while preserving the expected separation of GABA- and glycinergic ACs (**Fig. 6A**). Clustering in the integrated space identified 37 putative orthologous AC types, which we refer to as “non-mammalian oACs” (**Fig. 6B, Fig. S10E**). Notably, the correspondence was tight among the six well-sampled non-mammalian species (>3,000 cells), including lamprey, which diverged over 500 MYA (**Fig. 6B**). Using the sauropsids as a bridge between the two orthotype analyses, we found that 38 of the 42 mammalian oACs (88%) had a statistically significant match (*p* < 1×10^-6^, hypergeometric test) in the non-mammals and 26 of these (62%) were 1:1 correspondences (**Fig. 6C**). Each of these orthologies was supported by >40 homologous genes, demonstrating a high degree of molecular conservation (**Fig. 6D**). These included conserved signatures for oAC1 [A2], oAC2 [VG3], oAC6 [A8], oAC8 [nGnG-gl], oAC30 [PDGFRA], oAC36* [nGnG-ga], oAC37* [CRH], oAC42* [SAC] as well as many other less well studied oACs. A subset of these markers is highlighted in **Fig. 6E**. These findings provide strong evidence that AC diversity originated in basal vertebrates over 500 MYA.

**Fig. 6.**
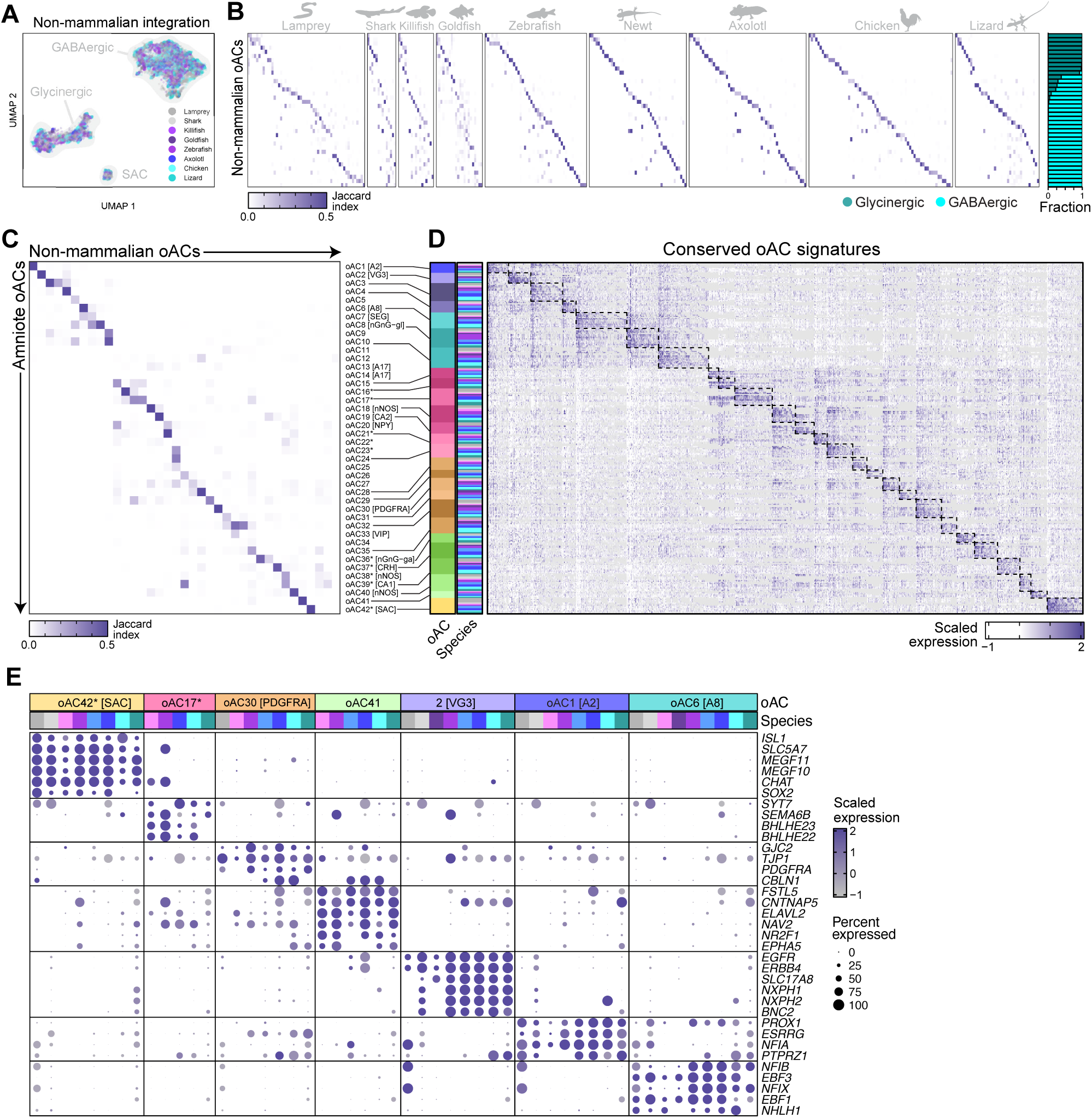
Non-mammalian oACs and their correspondence with amniote oACs. A) UMAP embedding of the SAMap integration of non-mammalian ACs. Cells are colored by species. Major clusters correspond to GABAergic ACs, glycinergic ACs, and SACs. B) Heatmaps showing the correspondence (Jaccard index) between unsupervised clusters from SAMap integration (rows) of non-mammalian ACs to the within-species clusters (columns). Rightmost bar graph shows the subclass composition (GABAergic vs. glycinergic) within each oAC. C) Correspondence (Jaccard index) between non-mammalian oACs (columns) and amniote oACs (rows), estimated by matching the oAC labels of chicken and lizard, which were common to both clusterings (see **Methods**). D) Heatmap showing the expression of top differentially expressed homologous gene combinations identified by SAMap for the oAC groups. Rows represent cells ordered by species within each nonmammalian oAC, and columns are differentially expressed homologous gene combinations. E) Dotplot showing markers (rows) for a subset of oACs conserved across vertebrates. Columns correspond to species types that mapped to a given oAC, while rows correspond to homologous gene combinations.

### Coordinated evolution of AC and RGC diversity

Although several AC types were highly conserved across species, we observed substantial variation in their overall diversity. Even among well-sampled species (>7k cells), the number of molecularly distinct AC clusters ranged from 32 types in marmoset to 66 types in chicken. We hypothesized that this variability in AC diversity may be linked to the diversity of RGCs (*78*), which constitute the output channels of the retina.

Across species with >3k cells profiled of either class, AC and RGC diversity were linearly correlated (R = 0.91, *p* < 0.01) with a slope ≈1.09 and intercept ≈17.6 (**Fig. 7A**). This relationship was not explained by sampling depth, as neither AC nor RGC diversity correlated with the number of cells profiled (**Fig. S11A**). This positive association could be recovered using an alternative measure of diversity (R = 0.89, *p* < 0.01) (**Fig. S11B**). By contrast, AC diversity did not correlate with BC diversity, nor did RGC diversity correlate with BC diversity (**Fig. S11C**), indicating a specific pattern of coordinate evolution between ACs and RGCs.

**Fig. 7.**
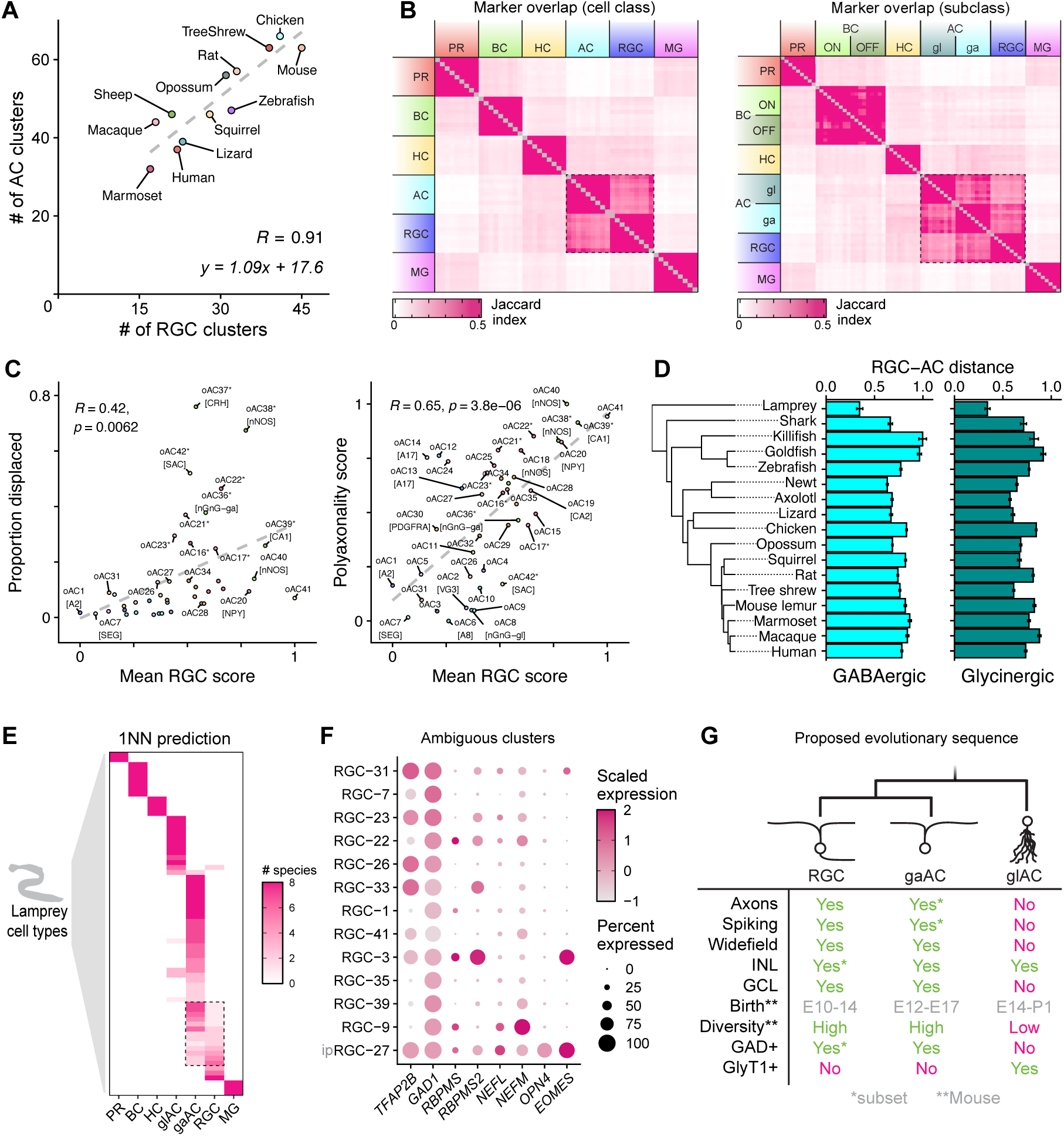
Shared evolutionary origins of GABAergic ACs and RGCs. A) Scatter plot of AC diversity vs RGC diversity across species. Pearson R = 0.91, *p* < 10^-4^ (correlation test). B) *Left:* Heatmap showing the marker overlap (Jaccard index, see **Methods**) between cell classes from the nine non-mammalian species (order from left to right: chicken, lizard, axolotl, newt, zebrafish, goldfish, killifish, shark, lamprey). ACs and RGCs are most similar. *Right:* Same as left, but with ACs split by GABA and glycinergic subclass. RGCs are more similar to GABAergic ACs than glycinergic ACs. C) *Left:* Scatter plot showing the relationship between proportion displaced (y-axis) vs RGC score (x-axis) across oACs. Frequently displaced ACs tend to have higher RGC scores. *Right:* Scatter plot showing the relationship between polyaxonality score (y-axis) vs RGC score (x-axis) across oACs (see **Supplementary Materials and Methods** for score details). The most RGC-like oACs are putative polyaxonal types. The correlation was preserved even when removing 11 axon components that overlapped with RGC module genes (R = 0.6, p < 10^-4^). D) Normalized RGC distance for GABAergic and glycineric ACs. Distance between ACs and RGCs is smallest in lamprey, the most basal vertebrate in our dataset. E) Heatmap showing the 1-nearest neighbor (1NN) prediction of lamprey clusters based on the SAMap alignment matrix from the non-mammalian cell class integration (**Fig. S10A**). Values in the heatmap represent the number of non-mammalian species that support each prediction (8 reference species total). Dashed box highlights 13 lamprey clusters that map ambiguously to both GABAergic ACs and RGCs. F) Dotplot showing expression of GABAergic AC and RGC markers in the 13 clusters from Fig. 7E. The numbering of the clusters is as in Ref. (*25*). Lamprey has at least two *RBPMS*^+^*GAD1*^+^*TFAP2B*^+^ clusters. One of these (RGC-27) is the putative ipRGC (*OPN4*^+^*EOMES*^+^). G) Table showing the morphological and physiological characteristics shared between GABAergic ACs and RGCs. Glycinergic ACs, in contrast, share few of these properties.

### Shared evolutionary origin of GABAergic ACs and RGCs

We next assessed the relationship of ACs and AC subclasses to other retinal cell classes. Based on marker overlap, ACs and RGCs are the most transcriptionally similar retinal cell classes across multiple species (**Fig.** Error! Reference source not found.**B**, left). Moreover, GABAergic ACs were more similar than glycinergic ACs to RGCs (**Fig.** Error! Reference source not found.**B**, right). Finally, within the GABAergic AC population, types most similar to RGCs were typically displaced and/or putative polyaxonal types, based on axon component genes (**Fig. 7C**). The implications of these observations are discussed below.

To assess whether the transcriptional difference between ACs and RGCs varies across evolution, we computed the distance between these two cell classes (averaged across types) in principal component space and normalized it to a reference distance (**Fig. 7D**, see **Methods**). This analysis revealed greater similarity between GABAergic ACs and RGCs in the most ancient vertebrates, lamprey (*p* < 0.01, β = −0.42, phylogenetic linear model) and shark (*p* < 0.05, β = −0.27, phylogenetic linear model) (**Fig. 7D**).

We therefore examined the lamprey retinal atlas, representing a lineage that diverged over 500 MYA (*25*). Using SAMap, we predicted cell class identities of lamprey clusters using the other eight non-mammalian species as reference. Consistent with the original study (*25*), lamprey clusters mapped unequivocally to PRs, BCs, HCs, glycinergic ACs, and MG in other species (**Fig. 7E**). In contrast, 13 clusters mapped to a similar extent to RGCs and GABAergic ACs. All 13 clusters expressed *GAD1,* and 9/13 expressed *TFAP2B,* consistent with GABAergic AC identity (**Fig. 7F**). However, 8/13 clusters also expressed *RBPMS* or *RBPMS2*, and all expressed the neurofilament genes *NEFL* and *NEFM*, consistent with RGC identity.

Notably, one ambiguous cluster corresponded to a putative intrinsically photosensitive RGC (*OPN4*^+^*EOMES*^+^), suggesting that ipRGC precursors may have been AC-like (**Fig. 7F**). Although *GAD1*/*2*+ RGCs have been observed in other species, these never expressed the AC transcription factor *TFAP2B*. In addition, several *RBPMS*+ clusters expressed *NPY* and *NOS1*, markers of GABAergic AC subpopulations in other vertebrates.

Together, these data support a model in which modern day ACs and RGCs arose from a common ancestral precursor (**Fig. 7G**). Glycinergic ACs diverged first, followed by a bifurcation giving rise to RGCs and GABAergic ACs. These two populations subsequently diverged further, with some GABAergic AC types such as displaced or polyaxonal ACs—likely diverging later than others. This latter split likely occurred around the cyclostome–gnathostome divergence (∼560 MYA), consistent with the presence of several lamprey clusters that map ambiguously to RGCs and GABAergic ACs of other vertebrates.

## Discussion

Our cross-species integration reveals that much of the diversity of AC types, the most heterogeneous neuronal class in the retina, is evolutionarily ancient and organized by a conserved regulatory logic among vertebrates. Here, we summarize our main results and the conclusions we draw from them.

### Evolutionary conservation of orthotypes

Despite extensive heterogeneity, integration across amniotes resolves 42 molecularly conserved AC orthotypes (oACs) (**Fig. 1D**), many of which can be traced back to fish (**Fig. 6D**). The preservation of several one-to-one correspondences across vertebrates supports the view that a large fraction of modern AC diversity derives from an ancient repertoire. Accordingly, many apparent “species-specific” clusters may reflect divergence in marker usage or abundance rather than the *de novo* emergence of novel AC types.

More than half of the amniote oACs could be identified in non-mammals with high confidence (Jaccard index > 0.3). These include eight glycinergic types (oAC1 [A2], oAC2 [VG3], oAC3, oAC4, oAC6 [A8], oAC8 [nGnG-gl], oAC9, oAC10) and 16 GABAergic types (oAC16*, oAC17*, oAC18 [nNOS], oAC19 [CA2], oAC21*, oAC24, oAC28, oAC29, oAC30 [PDGFRA], oAC32, oAC36* [nGnG-ga], oAC37* [CRH], oAC39* [CA1], oAC40 [nNOS], oAC41, oAC42* [SAC]). The fact that well-studied types like A2 and SAC are conserved in non-mammals is not surprising given the ethological importance of the rod pathway and direction selectivity, respectively (*24*, *25*). However, the fact that many lesser known types are highly conserved is notable. These may contribute to object motion sensitivity (VG3, CA2), gap junction regulation (CA1), and light adaptation (nNOS)—as well as many other unknown functions.

One interesting mapping is the additional cholinergic type in zebrafish (*79*), which is morphologically and functionally distinct from SACs. We found clusters corresponding to this AC type in our zebrafish and killifish datasets based on the expression of *BHLHE22* and *BHLHE23*. Although members of this non-mammalian oAC express some classic cholinergic markers like *CHAT* and *SLC18A3*, they differ from SACs in other ways (**Fig. 6E**), and align with oAC17* and oAC29 in mammals.

### Conservation of gene expression

Within the conserved cellular repertoire of oACs, evolution flexibly tunes gene programs. For instance, only a small set of neuropeptides (*VIP*, *TAC3*, *CRH*, *NPY*, *SST*) shows strong cross-species conservation, suggesting selective retention of specific modulatory functions (**Fig. S8A,B**). In contrast, many adhesion molecules and connectivity-associated genes (e.g. *DSCAM*, *DSCAML1*, *IGSF11*, *NXPH1*, *NXPH2*, *CBLN1*, *CBLN2*) are broadly conserved, consistent with strong purifying selection acting on circuit wiring (**Fig. S8A,B**). Conserved patterns of connexin expression further point to type-specific connectivity rules, with *GJD2* enriched in A2 ACs (*80*, *81*) and *GJC1* enriched in multiple GABAergic oACs (**Fig. S8A,B**).

### Differences in AC type abundance across species

A major axis of evolutionary change appears to be redistribution of conserved types. We identify primate enrichment of oAC30 [PDGFRA], particularly in the fovea, hinting at a potential role in high-acuity visual processing (**Fig. 3E**). We also observed several cell type frequency differences between the diurnal rodent (rhabdomys) and the closely related rat, mouse and peromyscus, which are nocturnal. First, rhabdomys had a higher proportion of OFF SACs than its nocturnal counterparts (**Fig. 4F**), consistent with their higher proportions of OFF bipolar cell circuitry (*82*). Second, A17 ACs (oAC13/14) were depleted in the rhabdomys compared to other rodents (**Fig. S4**, **S5**), possibly due to its reduced demand for scotopic vision.

### Developmental hierarchy

We hypothesize that the deep conservation of orthotypes results from conserved developmental programs. Our parsimony-based reconstruction recapitulates known lineage structure such asseparation *TCF4*-associated glycinergic signatures and *MEIS2*-associated GABAergic families(*14*). The *SATB2^+^EBF3^+^*module was selectively activated in SEG and nGnG-gl ACs, consistent with prior work(*40*). Upstream of *SATB2* and *EBF3* was *PRDM13* (**Fig. 6B**), which is known to be essential for generating glycinergic ACs, including EBF3+ ACs(*83*, *84*). Notably, our parsimony analysis reveals several TFs (*ZFHX3, ZFHX4*, *MEIS1, ST18, ZEB2, PBX1*, and *PBX3*) and TF modules (e.g. *PBX1*/*MEIS1*) not previously implicated in the specification of AC types.

Some oACs clustered by known stratification pattern, e.g. oAC38* [nNOS], oAC37* [CRH], and oAC13/14 [A17] stratify in S5, oAC19 [CA2], oAC36* [nGnG-ga] and oAC31 [nNOS] stratify in S3, and oAC30 [PDGFRA] and oAC33 [VIP] stratify in S4 (**Fig. 5A**). The tendency of developmentally related oACs to share IPL stratification patterns suggests coordination between fate programs and synaptic targeting.

### Evolutionary history of GABAergic ACs and RGCs

Across species, AC diversity scales tightly with RGC diversity (slope ≈1), an association not explained by sampling depth or mirrored by bipolar cells. This specificity supports coordinated evolution between ACs and RGCs, consistent with the idea that expanding the retinal output channels is accompanied by elaboration of inhibitory circuitry that shapes those channels. Moreover, RGCs are most transcriptomically similar to ACs, suggesting a shared ancestry, even though the former are excitatory and the latter are, with few exceptions, inhibitory.

A key result is that RGCs are more closely related to GABAergic than to glycinergic ACs and, within the GABAergic clade, particularly close to displaced and/or putative polyaxonal ACs. Lamprey offers an informative window into this relationship. In its retina, multiple clusters map ambiguously between RGCs and GABAergic ACs and co-express canonical markers of both populations (*TFAP2B*, *GAD1*, *RBPMS*, *RBPMS2*).

Within this evolutionary framework, the persistence of RGC-like morphological, functional, and molecular features in specific GABAergic AC types, which may seem puzzling at first, is naturally interpreted as a remnant of shared ancestry. The similarities are numerous including presence in the GCL, expression of GABAergic but not glycinergic makers (*85*), and relatively early birth (*86*) (**Fig. 7G**). More broadly, this model raises the possibility that some modern class boundaries arose through incremental acquisition of distinct molecular programs—such as axon guidance cues—rather than through abrupt lineage separation. To recount Dobzhansky’s famous dictum, “Nothing in biology makes sense except in the light of evolution.”

### A vertebrate retinal orthotype atlas

Together with our previous work (*21*, *87*), this completes a vertebrate retinal orthotype atlas. Altogether, we have identifed 5 oPRs, 2 oHCs, 12 oBCs, 21 oRGC, and 42 oACs, which represent a conserved set of orthologous types that likely date back to the mammalian ancestor. Most of these date back to even earlier, perhaps to the vertebrate ancestor. By outlining cellular relationships across species, this lexicon provides a resource that can support future morphological and physiological studies across the tree of life.

## Materials and Methods

### Single-nuclei RNA-sequencing

#### Axolotl and newt

Retinas were collected from 5 wild type axolotls between 4-8 months old, as well as 5-6 month-old wild type Iberian ribbed newts. Salamander retinas were dissected in cold amphibian PBS (PBS + 25% volume water) and snap-frozen by submerging in an isopentane bath cooled on liquid nitrogen. Nuclei were isolated according to the described protocol (PMID: 39240519) using 30 seconds of lysis incubation.

Isolated nuclei were resuspended in 1X Diluted Nuclei Buffer (10X Genomics; PN-2000207), and 8000-16,000 nuclei were loaded per reaction for downstream assays. Single-nucleus transcriptomes were generated using the Chromium Next GEM Single Cell Multiome ATAC + Gene Expression Kit (10X Genomics cat.

PN-1000285) following 10X Genomics protocol CG000338 Rev F. Libraries were sequenced at the Novogene Sequencing Core (Sacramento, CA & Beaverton, OR) on the NovaSeq X Plus instrument (Illumina). Read alignment and cell calling were performed using CellRanger-ARC (10X Genomics).

#### Rat and mouse lemur

We enriched RGCs and BCs by using fluorescence-activated nuclei sorting for the nuclear antigens NEUN/RBFOX3 and CHX10/VSX2 respectively. Frozen retinal tissue was dissociated with a Dounce homogenizer in NP-40–based lysis buffer containing Tris, CaCl₂, MgCl₂, NaCl, RNase inhibitor, and DNase I. The suspension was filtered (40 µm), nuclei were pelleted (500 rcf, 5 min), and resuspended in staining buffer. Nuclei were incubated with NEUN antibodies for 12 min at 4 °C, washed, and resuspended in sorting buffer followed by DAPI staining. NEUN⁺ and CHX10⁺ nuclei were isolated by FANS (BD FACSDiva v8.02) and washed again before being placed in 0.04% BSA/PBS at ∼1,000 nuclei/µl. Nuclear quality was confirmed microscopically prior to loading ∼8,000 nuclei per channel onto a 10x Chromium Single Cell Chip. Since AC types express RBFOX3 non-uniformly, the cell type proportions in the raw atlases are not reflective of their natural abundance. However, in un-enriched collections, AC subtype proportions closely matched immunostaining-based estimates (**Fig. 2F**), suggesting that the un-enriched collections provide a faithful representation of native AC diversity.

Single-nucleus RNA-seq libraries for rat and mouse lemur were prepared using the Chromium 3′ v3 or v3.1 chemistry (10x Genomics), as done previously (*21*). Briefly, isolated nuclei were encapsulated in Gel Beads-in-Emulsion, where they underwent lysis and barcoded reverse transcription to generate cDNA. The resulting cDNA was then amplified, enzymatically fragmented, and tagged with 5′ adapters and sample indices to produce sequencing-ready libraries. Libraries were run on an Illumina NovaSeq at the Harvard Bauer Core Facility. Demultiplexing and alignment were performed with Cell Ranger (v4.0.0, 10x Genomics).

#### Squirrel and sheep

To obtain more amacrine cells from squirrel and sheep, we sequenced two additional NEUN-enriched channels for squirrel and two NEUN-enriched channels for sheep. We used the same protocol as for the rat and mouse lemur (see above).

All animal procedures were conducted in accordance with institutional guidelines and approved by the Institutional Animal Care and Use Committee (IACUC).

### Alignment of sc/snRNA-seq data

For newly sequenced species (mouse lemur, rat, axolotl, and newt), we aligned reads to the reference genome available on either ENSEMBL or NCBI. Mouse lemur and rat reads were aligned to the ENSEMBL genomes Mmur_3.0 and mRatBN7.2, respectively. Axolotl and newt reads were aligned to the NCBI refseq assemblies (GCF_040938575.1 and GCF_031143425.1, respectively). Although axolotl and newt RNA profiles were acquired using 10x Chromium Single Cell Multiome ATAC + Gene Expression, we did not use the chromatin accessibility data in this study. Further details regarding each atlas used in the study are in **Table S2**.

### Dimensionality reduction and clustering within each species

Expression matrices were loaded and analyzed using Seurat (v4.3.0). Briefly, counts from different cells were normalized by the total UMI count of each cell and log-transformed with a pseudocount of 1. The top 2000 highly variable genes were selected using the variance stabilizing transformation. These features were scaled across cells and PCA was run to reduce the dimensionality to 20-30 principal components. Graph-based clustering was performed in PC space using cosine distance. Uniform manifold approximate projection (UMAP) was performed on PCs to visualize the data in two dimensions. We used Doublet Finder with default parameters to detect and remove doublets (two cells in the same droplet). These were often easy to detect as they had a mixture of two very distinct signatures.

### Identification of gene homologs

To identify homologous genes, we used a reciprocal protein BLAST approach. To do so at scale, we used DIAMOND (*88*), a BLAST-compatible local alignment tool that runs 80-100x faster than BLAST. Protein sequences were downloaded from either NCBI or ENSEMBL for the same assembly used for aligning single-cell reads. Protein sequences were compared pairwise for similarity. Each protein sequence from species A was BLASTed against the full set of protein sequences from species B, and vice versa. Bit scores above an e-value < 1×10^-6^ were used as scores to determine 1:1 orthologs for Seurat integration. These scores were also used to initialize the SAMap homology graph.

### Integration of amniote atlases based on 1:1 orthologs

To integrate ACs across mammals, we used the Seurat CCA pipeline (*35*). We first subsetted each count matrix to 6,305 1:1 orthologous genes, then merge the Seurat objects. Cell types were downsampled to 100 cells to balance lowly and highly abundant cell types. Finally, Seurat (v4.3.0) CCA integration with 2000 variable genes and 20 principal components was used to integrate cells from different species into a common latent space. Cell type identities were not supplied at any time during this integration procedure. Graph-based clustering was performed in the latent space to define orthotypes (oACs). oACs were defined first for amniotes, and then for the non-mammalian vertebrates.

To compare amniote orthotypes to non-mammalian orthotypes, we used the common sauropsid species (chicken and lizard) as a bridge. The amniote and non-mammalian orthotype labels for chicken and lizard were mapped based on cell overlap, quantified by the Jaccard index (**Fig. 6C**).

To calculate marker-overlap between different non-mammalian cell classes/subclasses (**Fig. 7B**), we first integrated the atlases using Seurat CCA, and then found the top 500 differentially expressed genes for each cell class/subclass using the integrated assay to correct for species-level batch effects. We then computed the overlap (Jaccard index) of marker gene sets across cell classes/subclasses.

### Assessing robustness of amniote orthotypes

We applied several metrics to confirm the validity of the amniote oACs. To quantify the quality of the integration, we used the integration local inverse Simpson index (iLISI) and cell type local inverse Simpson index (cLISI). These were computed using the lisi R package (*36*). These metrics measure the degree of mixing of cells from different species (iLISI) and different cell types (cLISI). An ideal integration would have high mixing of cells from different species (high iLISI), but low mixing of cells from different types (low cLISI). To estimate the cLISI, we used our independently manually annotated SAC, A2, and VG3 cells. To confirm the accuracy of our oAC clustering, we computed precision (the ratio of true positives to the sum of true positives and false positives) and recall (the ratio of true positives to the sum of true positives and false negatives). Precision and recall values were computed for each species and for each manually annotated cell type (SAC, A2, VG3). These metrics could only be calculated for the SAC, A2, and VG3 oAC clusters, as there was no secondary, independent way to validate the inferred annotation of other oACs.

We also performed a resampling-based procedure to determine the robustness of the oACs. In each permutation, we randomly sampled 80% of cells and repeated the integration with randomly selected parameter values (principal components, k-nearest neighbors, and clustering resolution). Parameter values were randomly selected from a uniform distribution of values: 15-30 PCs, 16-24 k-nearest neighbors, and 1-2 resolution (spaced by 0.1). A total of 50 permutations were performed. Preservation of the original oAC clustering was observed visually by mapping the new clusters to the original clusters and coloring them by their identity (**Fig. S2D**). Streaks of the same color across permutations indicated that all oACs were highly robust, with some small oACs (e.g. oAC26 and oAC27) occasionally being merged in some permutations.

### Controlling for erroneous cross-species integration

In addition to the checks discussed above, we performed six control experiments to confirm that our cross-species integration pipeline has a low error rate and does not introduce artifacts:

1. We performed an integration of mouse BCs from Shekhar et al. (2016) with the rat retinal atlas. Mouse BCs integrated with rat BCs and did not intermix with other classes (error rate = 0.07%).
2. We integrated the mouse lemur atlas lacking BCs and rat atlas lacking RGCs. Mouse lemur RGCs and rat BCs remained distinct (0.02% error for rat, 1.1% error for mouse lemur).
3. We integrated amacrine cells from 17 amniotes where the expression values within species were randomly shuffled. While orthotypes corresponded to species-specific types in the real data (all ARI > 0.4), the orthotypes calculated from shuffled data had no correspondence to types calculated from shuffled data. This negative control indicates that the integration procedure does not artificially produce cross-species correspondence when there is none (all ARI < 0.01).
4. We integrated amacrine cells from 17 amniotes where the gene orthology key was randomly shuffled. Again, the integration algorithm produce an artificial correspondence (ARI = 0.2 for the integrated vs. original types in the species used as reference, but ARI < 0.07 in all others).
5. We integrated primate ACs where the expression signature of one type had been contaminated by another. Cell types mixed only when a substantial fraction (>= 0.5) of their signature was artificially perturbed.
6. We integrated mutually exclusive subsets of AC types. The resulting orthotypes were very similar to the original oACs (ARI = 0.64). However, three pairs of transcriptomically related oACs were merged erroneously across these subsets, representing an error rate of 6/42 (14.2%). The occasional merging of transcriptionally proximal types is an unavoidable issue, but can be mitigated with larger sample size or a targeted analysis of these subsets.

### Transferring metadata onto oACs

#### Proportion of displaced somata

The proportion of displaced somata for each mouse AC type from Yan et al., 2020 (*14*) was extracted from **Fig. 4C** of Choi et al. 2023 (*64*) using the Automeris WebPlotDigitizer. We transferred this measure onto the oACs, thereby inferring the displaced proportion for each orthotype, as a weighted average of the displaced proportion of its constituent mouse types, defined as:

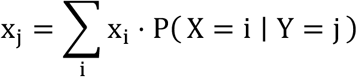

where *x_i_* is the proportion displaced for type *i* and P( X = i ∣ Y = j) is the fraction of orthotype *j* mapping to type *i*. This formula is exactly the conditional expectation 𝔼[*X* ∣ *Y* = *j*], and represents the optimal mean-squared-error estimator of the displaced proportion of orthotype *j*. The same formula was used to compute GABAergic and glycinergic scores for oACs (**Fig. 2A, C**), where the score (*x*_i_) within each species is the average expression of GAD1/2 (for GABAergic) or SLC6A9 (for glycinergic) in each species type, scaled to be between 0 and 1. Thus, GABAergic and glycinergic scores represent the average expression of these markers in each oAC, weighted by the contribution of their constituent types.

#### Frequency

The frequency of each orthotype could not be computed directly because species types were downsampled to 100 cells each prior to integration. Thus, we estimated this value indirectly using a weighted sum. We defined the frequency of orthotype j in a given species as the sum of the frequency of its constituent species types, weighted by the strength of their mapping:

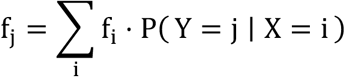

where *f_i_* is the frequency of species type i and P( Y = j ∣ X = i) is the proportion of type i mapping to orthotype j. Unlike the proportion displacement score, this frequency score has an additive property such that if an orthotype lumps together multiple species types, it will sum each of their individual frequencies. The frequencies shown in **Fig. 4** and **Fig. S7** are at the species level. In **Fig. S4** and **S5**, this analysis was done at the replicate level. To avoid estimating frequencies in species where an oAC may be absent, frequencies for each oAC were only estimated in species where that oAC was robustly detected (FDR < 0.10).

### Immunohistochemistry

We used banked retinal tissue for histology in human, macaque, mouse, guinea pig, and rabbit. All samples had been fixed in 4% paraformaldehyde for 30 minutes at 22°C. Samples were cryoprotected in graded sucrose solutions (10%, 20%, 30%) and stored at −20°C until use. For immunostaining, retinas were washed in phosphate-buffered saline (PBS) for 15 minutes, then blocked for 1 hour in 10% normal horse serum (NHS), 1% Triton X-100, and 0.025% NaN_3_ in PBS at 22°C. Primary antibodies were diluted in 3% NHS, 1% Triton X-100, 0.025% NaN_3_ and applied overnight at 22°C. The samples were washed with PBS, then incubated overnight with secondary antibodies diluted in 3% NHS, 0.1% Triton X-100, 0.025% NaN_3_. The samples were washed three times and counterstained with Hoechst 33342. The samples were washed three more times and a coverslip was mounted with Mowiol.

For imaging, confocal laser scanning images were acquired on a Zeiss LSM 880 microscope with a Zeiss Plan-Apochromat 20x/0.8 air DIC objective. We used an excitation wavelength of 488 nm for Alexa-488, 594 nm for Alexa-594, and 633 nm for Alexa-647. Adjustments to brightness, contrast and pseudo-color were made in FIJI (*89*).

### Analysis of ON and OFF starburst amacrine cells (SACs)

We also used the Seurat CCA pipeline to integrate starburst amacrine cells (SACs) across different clades – primates, rodents, laurasiatherians, sauropsids, and amphibians (**Fig. 4C**). In most species, the SACs cleanly split into two clusters. Within each clade, we clustered the integrated data at multiple resolutions and identified the resolution that gave two clusters. These clusters were then annotated into ON and OFF based on the expression of known ON and OFF markers identified previously in mouse and lamprey (*25*, *49*). We were unable to identify ON and OFF SACs in the four primates (human, macaque, marmoset, mouse lemur) because SACs formed only one homogenous cluster. SACs were too sparse in pig and ferret to perform this analysis. No SACs were identified in goldfish. We found SACs in killifish and catshark, but they could not be subclustered.

In an attempt to recover ON and OFF SACs in primates, we tried two supervised classification approaches: 1) a gradient-boosted tree classifier (XGBoost) trained on mouse ON and OFF SACs from Peng et al. 2020, and 2) linear discriminant analysis trained using the closest species that was successfully subclustered. Neither method generated trustworthy classifications, and we did not pursue this further.

### Regulatory tree construction by maximum parsimony

We sought to construct an oAC regulatory tree based on discrete transcription factor (TF) expression patterns using phylogenetic methods. First, we identified 87 TFs with conservation scores greater than 0.3 (see Supplementary Materials & Methods for definition). Then, we binarized the expression of each TF using k-means clustering, which generated more reliable results than Gaussian mixture models. We retained 70 TFs whose binarization explained at least 65% of the original expression variance, and confirmed the quality of binarization by visual inspection. We generated a TF by oAC matrix (70 x 42), on which we applied four phylogenetic clustering algorithms (maximum parsimony, UPGMA, maximum likelihood, and neighbor joining). We decided to use maximum parsimony (MP) because it can reconstruct intermediate states and, moreover, it exhibited greater stability than maximum likelihood, the only other algorithm able to reconstruct intermediate states. MP trees were generated using the “pratchet” function from the package phangorn (*66*) (v2.12.1) with 1000 minimum iterations and returning all equally parsimonious trees. This yielded 132 trees, using which we constructed a majority-rule consensus tree (**Fig. 5A**, top).

To confirm the robustness and validity of the consensus tree, we computed 70 leave-one-TF-out trees for each of the four methods (**Fig. 5A**, heatmap). We then checked whether clades suggested by the consensus MP tree were preserved in the 280 trees across all four methods. We did this for a set of coarse-resolution clades and a set of fine-resolution clades. In general, clades were supported by at least three of the four tree construction methods.

For reconstructing intermediate states, we determined the position of character changes using the anc_pars function, which relies on the ACCTRAN criterion (*67*). By this criterion, ambiguous character-state changes are assigned as early as possible on the phylogeny, favoring reversals over parallel gains. Branch lengths of this tree represent the number of TF switches (**Fig. S9**). Finally, these TF gain and loss events were projected onto the tree. We note that multiple equally parsimonious reconstructions exist; therefore, the tree presented in **Fig. S9** should be considered one potential reconstruction, yet to be validated. Analysis of protein-protein interactions is described in the **Statistical Methods** section.

### Integration of non-mammalian atlases

To account for the many-to-one gene homology pattern prevalent among distantly related species, we used SAMap (*77*) to integrate atlases across the non-mammals (chicken, lizard, axolotl, newt, zebrafish, killifish, goldfish, shark, lamprey). The same analytical pipeline was used for the cell class level integration, the bipolar cell type integration, and the amacrine cell type integration. Briefly, raw counts from Seurat were converted to h5ad files using SeuratDisk. Then, a custom SAMap script was used to downsample each cluster to 100 cells, run the SAM algorithm on each object, and then integrate the different species using SAMap run. To generate orthotypes for the non-mammals, we used Leiden clustering (*90*) on the stitched cell-by-cell graph generated by SAMap. We tried several clustering resolutions and multiple subsamples of cells to ensure that our downstream conclusions were robust across parameter choices. The resulting homology patterns were visualized using Jaccard index heatmaps as done for the amniotes.

With this pipeline, we performed three integrations: 1) all six cell classes (photoreceptors, bipolar cells, horizontal cells, amacrine cells, retinal ganglion cells, and muller glia); 2) bipolar cells; and 3) amacrine cells. The first two were used primarily to confirm that SAMap recovered the expected homology patterns, while the amacrine cell integration was used to infer AC type conservation in non-mammals. The six-class integration was also used for the one-nearest neighbor classification of lamprey cell types (**Fig. 7E**). For each lamprey cluster, we found its nearest neighboring cluster in the other 8 non-mammals based on the alignment matrix from SAMap. The cell class identity of the nearest neighbor was used to predict the identity of the lamprey cluster. Alignments with scores less than 0.05 were considered unmapped to avoid low-confidence predictions.

### Quantification of AC and RGC diversity

To determine the relationship between AC and RGC diversity, we subsetted to the 14 species in which there were at least 3,000 ACs and 3,000 RGCs. We further removed peromyscus and rhabdomys as the number of AC clusters had not saturated in these two rodents, suggestive of undersampling (**Fig. S11E**). Thus, we were left with 12 species for the correlation analysis. We first related the number of clusters of ACs, RGCs, and BCs to the number of cells sampled. No significant relationship was observed (all Pearson R < 0.3, all *p* > 0.4), suggesting that in these atlases, the number of clusters was independent of sampling bias. We then checked the relationship between the number of clusters in all three pairwise comparisons (AC vs RGC, AC vs BC, RGC vs BC). Only the AC vs RGC comparison was significant. To check the reproducibility of this result, we also repeated the analysis using the inverse Simpson diversity index, which is more robust to the choice of clustering resolution. The inverse Simpson index was computed using the cluster labels as:

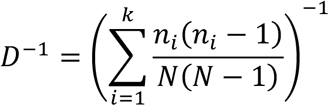

where *k* is the number of clusters, *n*_*i*_ is the number of cells from cluster *i*, and *N* is the total number of cells. The value *D*^-1^ represents the effective number of clusters, since if all clusters are equally represented, *D*^-1^ approaches *k*. Using this metric, we also found a signficant correlation between AC and RGC diversity, but not for the other two comparisons.

### Statistical Analysis

Statistical analysis was performed in R (v4.4.1). Correlation significance was determined using the cor.test function to compute *p*-values, with values less than 0.05 being considered significant. P-values were adjusted for multiple hypotheses using the Benjamini-Hochberg procedure.

#### Significance level for presence scores

To determine whether an oAC was present in a species, we first established a null distribution of presence scores by iteratively removing one of the well-characterized AC types (A2, SAC, VG3) from a given species, repeating the integration, and calculating its presence score (**Fig. S7A**). The median presence score for a missing cell type was 0.01 (**Fig. S7B**), indicating that null presence scores were generally quite low. Next, we determined the cutoff for a false discovery rate (FDR) of 0.10 by finding the threshold at which false positives over all positives was 0.10. This threshold was similar to a *p*-value threshold of 0.10; thus, our choice of FDR over *p*-value does not change the outcome significantly.

#### Enrichment of protein-protein interactions

Protein-protein interactions were downloaded from the STRING database (v12) accessed using the STRINGdb R package (*73*). We considered all pairs of genes residing in the same node of the tree as potential interactors, and identified 43 pairs that had a significant interaction score (>400). To generate a null distribution for the number of interactors expected by random placement, we then shuffled the position of genes across the nodes of the tree 10,000 times. Only 47 out of 10,000 permutations yielded at least 43 interactors, representing an observed p-value of 0.0048. Permutation *p*-value was computed as (b+1)/(N+1), where b is the number of permuted statistics at least as extreme as the observed statistic and N is the number of permutations.

#### Divergence analysis

We computed distances between AC and RGC cell types in a 3-dimensional (PC) space within each species. Three PCs were chosen as the dimensionality of the pseudo-bulked expression space (2000 top highly variable genes) was between 2-4 PCs in most species. We used 1000 bootstraps to compute 95% confidence intervals for the AC–RGC distance. We computed the distances separately for GABAergic and glycinergic ACs.

Distances were normalized to a reference distance (the average GABAergic AC–BC distance) to make them comparable across species. Finally, we used phylogenetic linear models (*91*) to determine whether the difference was significant between lamprey—or lamprey and shark—versus the remaining species. We used Pagel’s λ model to account for the phylogenetic covariance structure.

## Acknowledgments

Axolotls were obtained from the Ambystoma Genetic Stock Center (AGSC; University of Kentucky). Computational analysis was performed on the Savio cluster hosted by Berkeley Research Computing. We thank Chris Paciorek, Mark Yashar, and Sapana Soni for their computing advice and support. Shared microscopy equipment was supported by the UC Berkeley Vision Science core grant P30EY003176. Confocal imaging was conducted at the CRL Molecular Imaging Center (RRID: SCR_017852), supported by the Helen Wills Neuroscience Institute. We thank Holly Aaron, Luis Alvarez, and Feather Ives for their microscopy advice and support. We thank Greg Field for helpful discussions on designing control experiments for cross-species integration and Greg Schwartz for helpful discussion regarding displaced amacrine cells. We thank Marla Feller, Tom Baden, and George Kafetzis for feedback on the manuscript.

## Funding

National Institutes of Health grant EY028625 (KS)

National Institutes of Health grant EY024265 (TP)

National Institutes of Health grant F31EY038101 (DT)

National Science Foundation CRCNS grant 2309039 (KS, DT)

National Institutes of Health grant MH105960 (JRS)

National Institutes of Health grant EY028633 (JRS)

McKnight Foundation (KS)

Glaucoma Research Foundation (KS) BrightFocus Foundation (KS)

Research Corporation for Science Advancement #SA-MND-2023-031b (KS)

ERC grant DEEPRETINA (101045253) (OM)

## Author contributions

Conceptualization: DT, JRS, and KS;

Software: DT, VD, and JH;

Formal analysis: DT, VD, and JH;

Investigation: DT, AM, JT, OM, SB, and TP;

Writing – original draft: DT, JRS, and KS;

Writing – review and editing: DT, JRS, and KS.; Supervision: JRS and KS

Funding acquisition: JRS and KS

## Competing interests

Authors declare that they have no competing interests.

## Data and materials availability

All data are available in the main text or the supplementary materials. The raw and processed sequencing data produced in this work are available via the Gene Expression Omnibus (GEO) under accession GSE237215. The accession numbers for species-specific datasets are listed in **Table S2**, including for previously published data. Species reference genomes are available on Ensembl (https://www.ensembl.org) or NCBI (https://www.ncbi.nlm.nih.gov/datasets/genome/). Code to reproduce the analyses is available on GitHub (https://github.com/shekharlab/AmacrineCells). Interactive visualization of orthotype atlases is available on the Single Cell Portal (https://singlecell.broadinstitute.org/single_cell).

## Supplementary Text

### Identification of oAC-specific markers

To identify genes that specifically label an oAC, we derived a specificity score inspired by convolutional filters. For each gene, we consider a 17 species x 42 oAC matrix, where each value represents its expression in that species/oAC combination. An ideal marker would be expressed (value = 1) in one column (oAC) in all 17 species, but not expressed elsewhere (value = 0). To construct the observed matrices, we took the vector of normalized counts and scaled expression for each gene across oACs (within each species separately). We then concatenated these 42-length vectors across species, building the 17 species x 42 oAC matrix for each gene. We refer to this matrix as *M*. Each matrix was normalized such that the maximum possible z-score equals 1. We then subtract this matrix from the idealized matrix for each oAC (a matrix with an entire column of 1s for that oAC, and 0s in every other cell) and compute its norm (L1-norm):

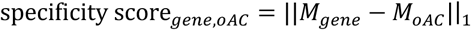

A score is computed for each gene/oAC combination. Since lower values represent more specific markers, we centered this value around the emprically computed random score (determined by shuffling expression values across species) and scaled it by the best possible score (score = 0). Using this normalized specificity score, a score of 1 indicates a perfectly specific oAC marker and a score of 0 indicates a random marker. Note that scores can be slightly negative.

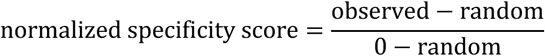

### Identification of conserved genes

To identify genes whose expression is conserved across species, we derived a conservation score. The conservation score is simply the mean Pearson correlation between the rows of the expression matrix *M* (17 species x 42 oAC) described above. If the expression in one species is informative of expression in other species, than the score will be near 1. If the expression in one species is not informative of expression in other species, the score will be near zero. No further normalization was applied to this score.

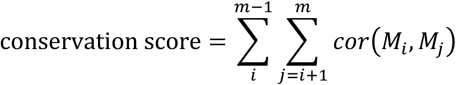

where *M_i_* is the ith row of *M*.

### RGC and polyaxonality module scores

For module scores, we used the AddModuleScore function from Seurat, which computes a single expression enrichment score in each cell for a given gene set relative to a randomly selected set of genes (matched for expression level). For the polyaxonality score, we used 338 genes in the axon component category (GO:0030424) of the gene ontology database from NCBI (https://ftp.ncbi.nlm.nih.gov/gene/DATA/gene2go.gz). For the RGC module, we used the top 50 differentially expressed genes from the RGCs in the 17 species integration from Hahn et al., 2023. Scores were min-max normalized across oACs, such that 0 represents lowest score and 1 the highest score.

---

**Fig. S1.**
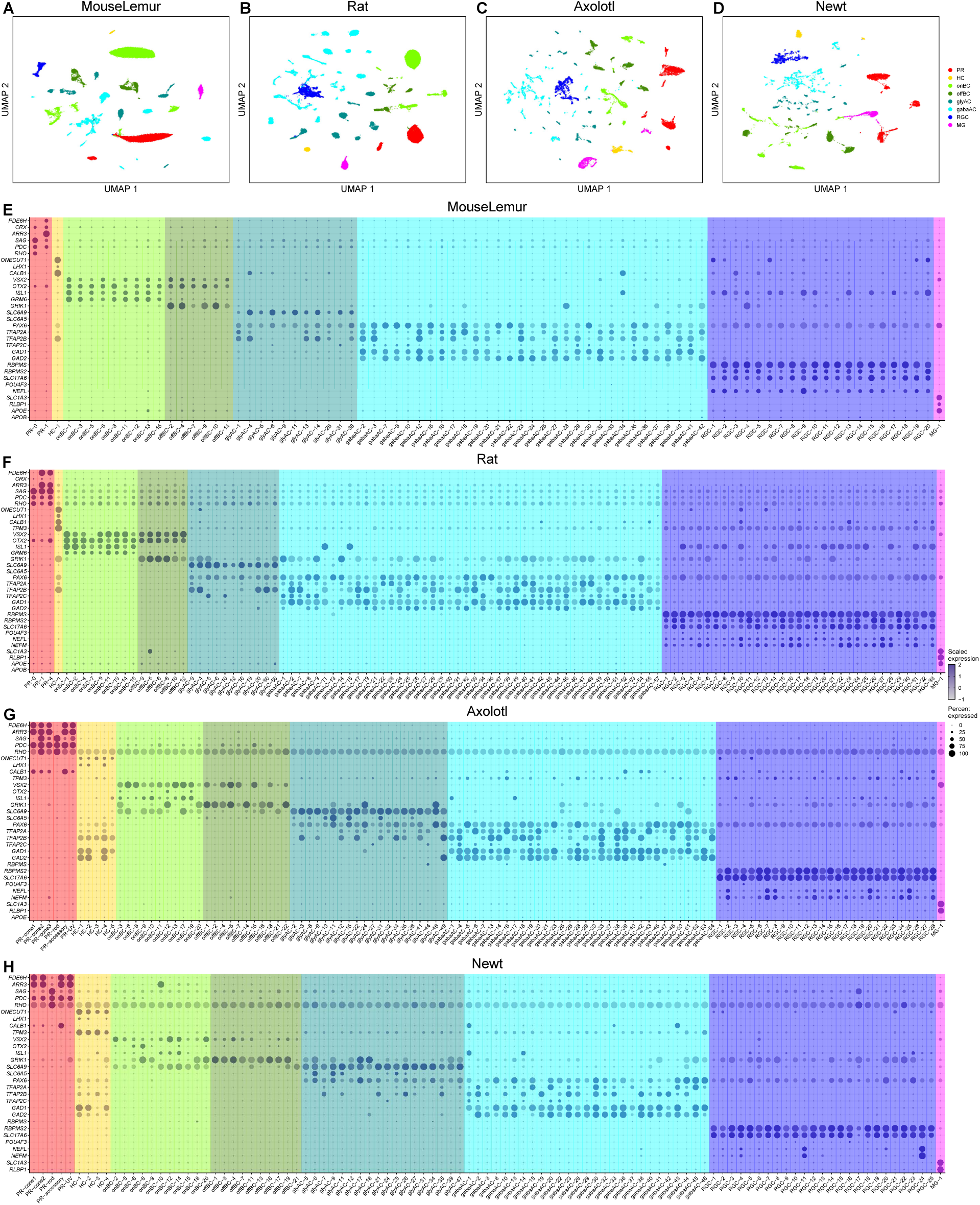
Retinal atlases for mouse lemur, rat, axolotl, and newt. A-D) UMAP embeddings for mouse lemur (A), rat (B), axolotl (C), and newt (D). Cells colored by class identity. Red: photoreceptors (PR); light green: ON bipolar cells; dark green: OFF bipolar cells; yellow: horizontal cells; dark cyan: glycinergic ACs; cyan: GABAergic ACs; blue: RGCs; magenta: Müller glia. E-H) Dotplots showing the expression patterns of cell class marker genes (rows) in transcriptionally defined clusters (columns). Clusters are grouped by class identity as in panels A-D.

**Figure S2.**
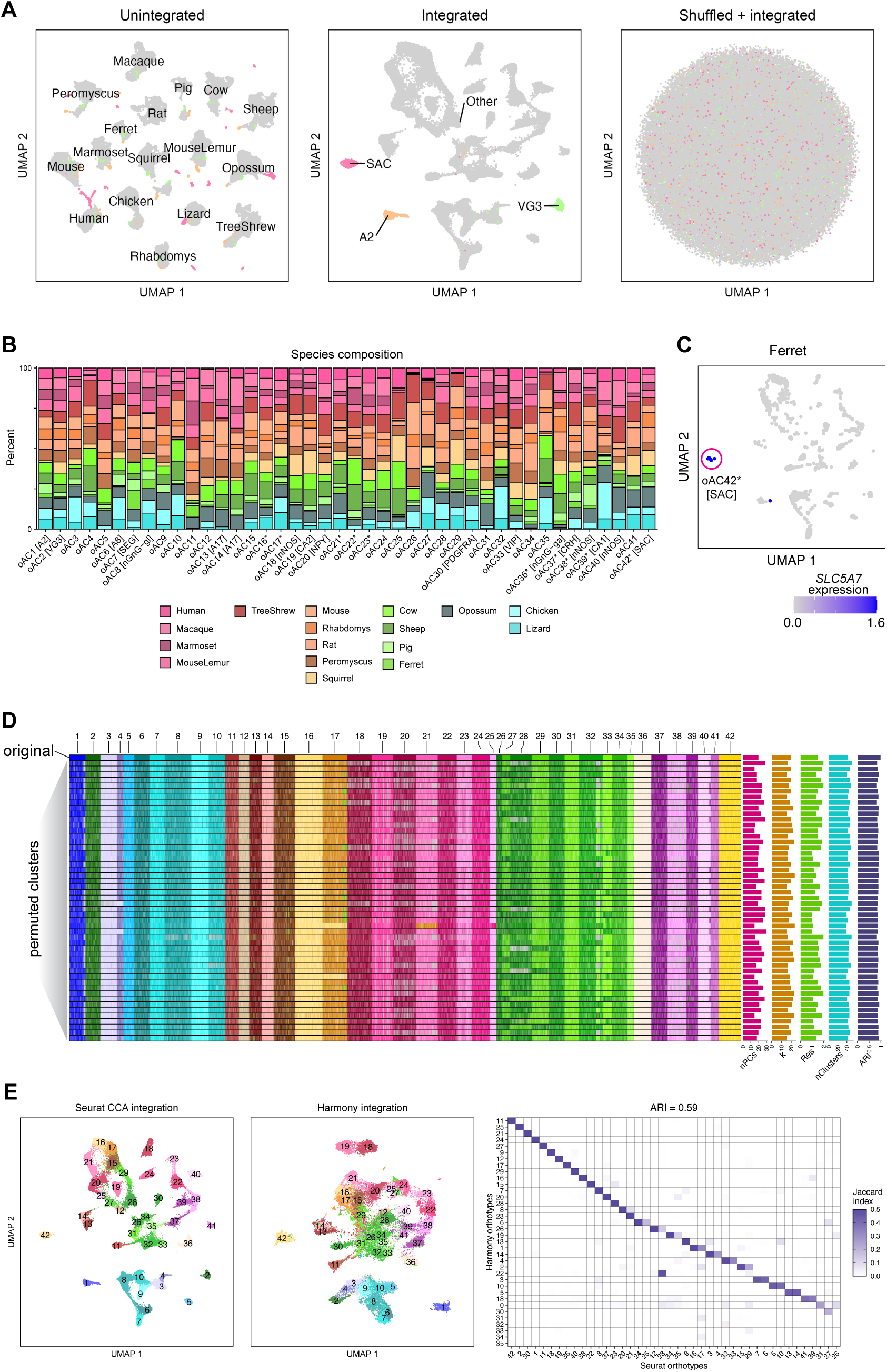
Integration of mammals and robustness analysis. A) UMAPs showing the unintegrated (left), integrated (middle), and shuffled + integrated cells (right). Notice that manually annotated orthologous types (SAC, A2, VG3) cluster together only in the middle panel corresponding to the integrated embedding. B) Stacked bar graph showing the species composition of each oAC from **Fig. 1D.** C) Feature plot showing the expression of *SLC5A7* in ferret ACs. The integration identifies 7 putatively true SACs in ferret, which were too sparse to form a distinct cluster in the ferret data. However, these cells co-cluster with other SACs in the integrated dataset. D) Heatmap showing the robustness of oACs to various choices of parameters. Each column represents a cell and each row represents a clustering of the cells. The top row shows the oAC clustering, and each row below shows the permuted clustering (n = 50 permutations). The oACs are colored by their best matching oAC from the original integration. A cell is given the color grey if its cluster in that permutation did not have a matching oAC. A cell is given the color white if it was removed due to subsampling. The fact that oACs form colored columns indicates that they are robust to different choices of parameters. E) UMAPs comparing the embedding space from Seurat integration (left) to Harmony integration (middle). Confusion matrix on the right shows a good correspondence between Seurat clusters and Harmony clusters.

**Figure S3.**
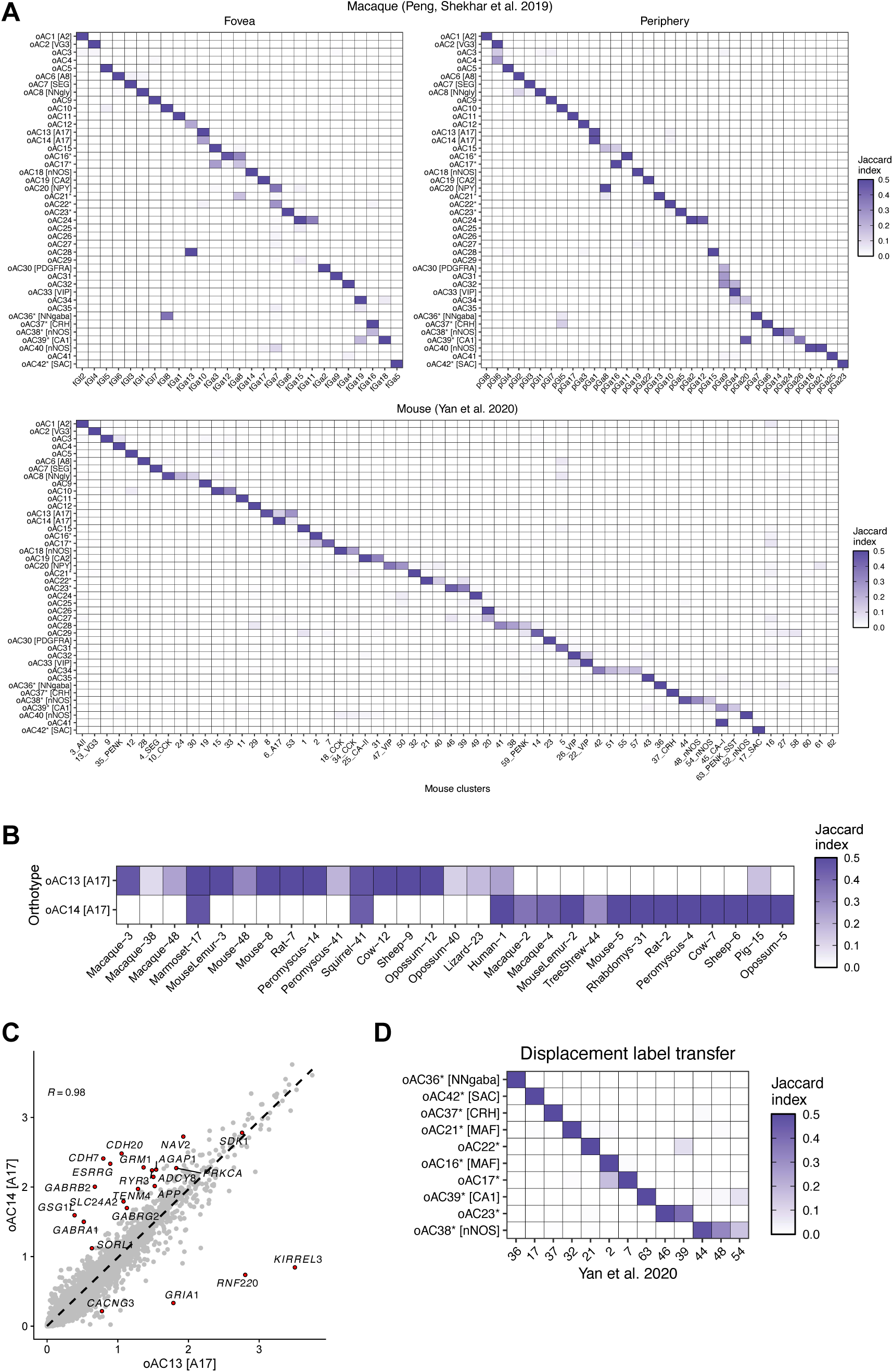
Meta-analysis of orthotypes. A) Confusion matrix showing correspondence between oACs and AC types from macaque fovea and periphery (Peng, Shekhar et al. 2019; top two panels) and mouse (Yan et al. 2020; bottom panel) atlases. oACs display a specific correspondence with species-specific types B) oAC13 and oAC14, both of which map to A17 ACs, map to distinct clusters within most species. This suggest that they may each represent a genuine cell type. C) A17-like oACs 13 and 14 are transcriptionally similar but distinguished by several functionally important genes (e.g. AMPA receptors, *GRIA1* and the adhesion molecule Kirrel3). D) The 13 mouse types containing displaced cells (based MERFISH atlas in Ref. 13) converge onto ten oACs, all of which are GABAergic. We refer to these oACs as “displaced oACs”.

**Figure S4.**
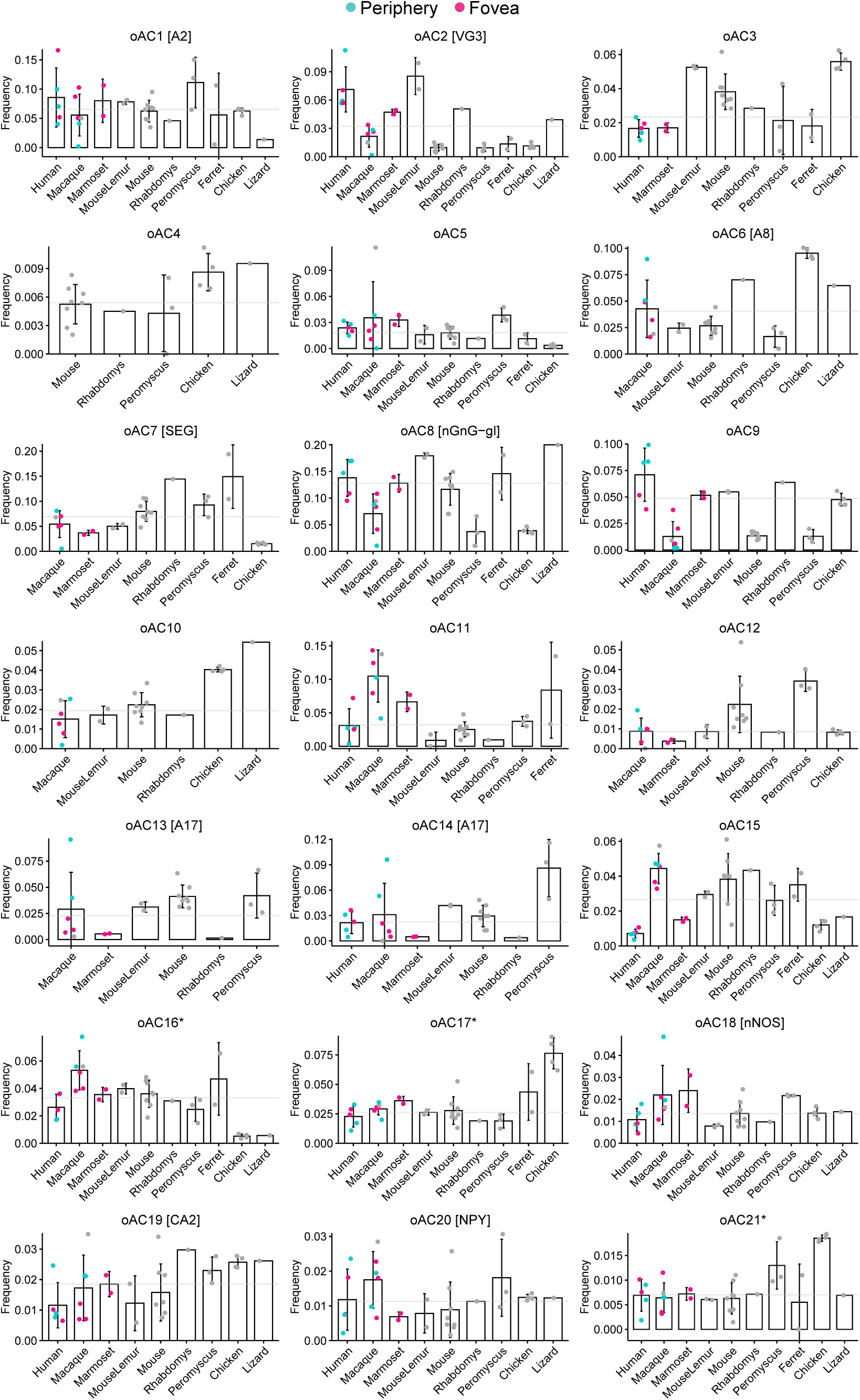
Cell type proportions quantified from unenriched batches (oACs1-21). Each bar graph shows one oAC. For higher primates, we show both foveal (pink) and peripheral (orange) collections. Only unenriched batches were used for quantification; hence, we have few species for some orthotypes. Grey line shows the median frequency across all species.

**Figure S5.**
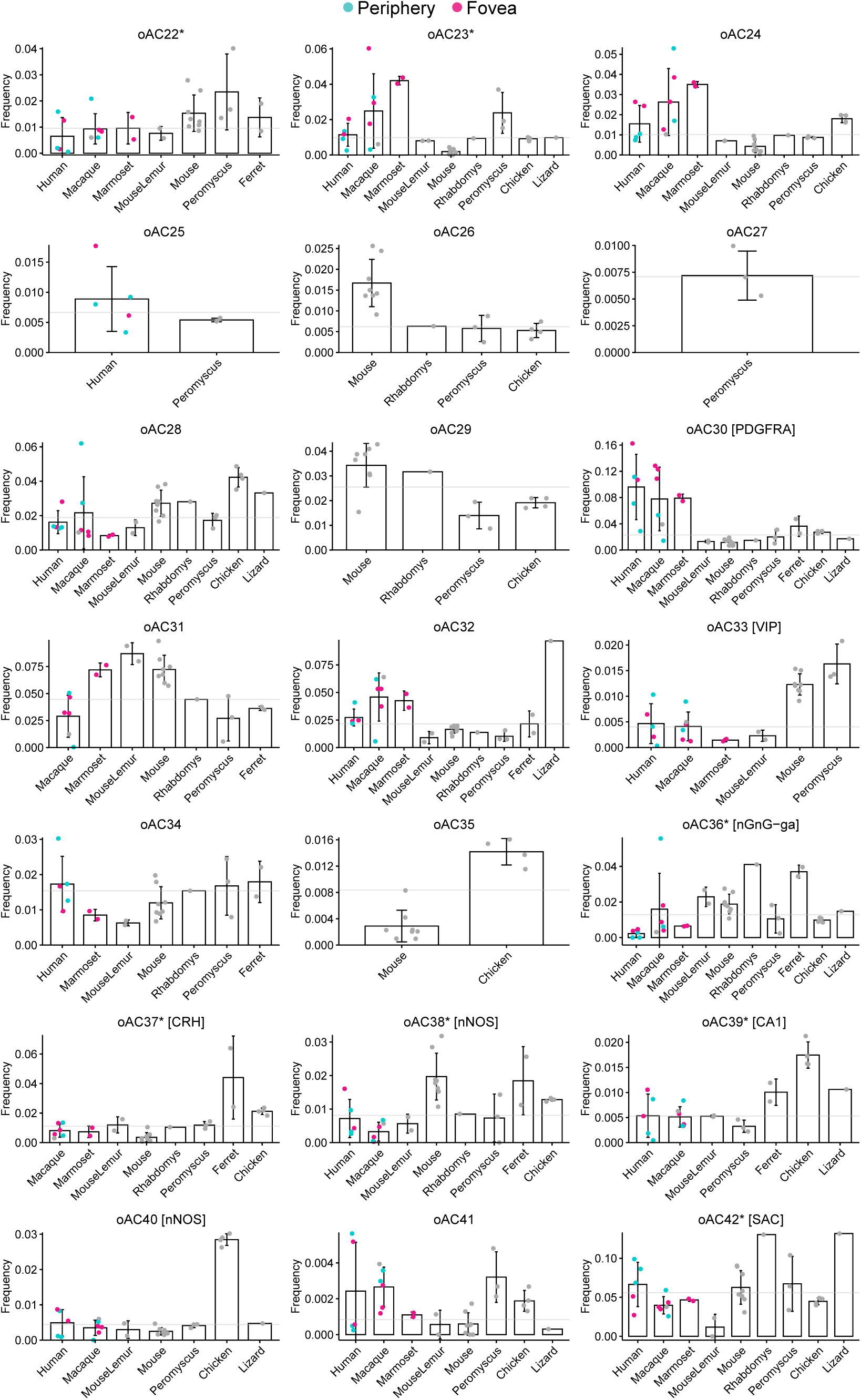
Cell type proportions quantified from unenriched batches (oACs22-42). See **Fig. S4** for details.

**Figure S6.**
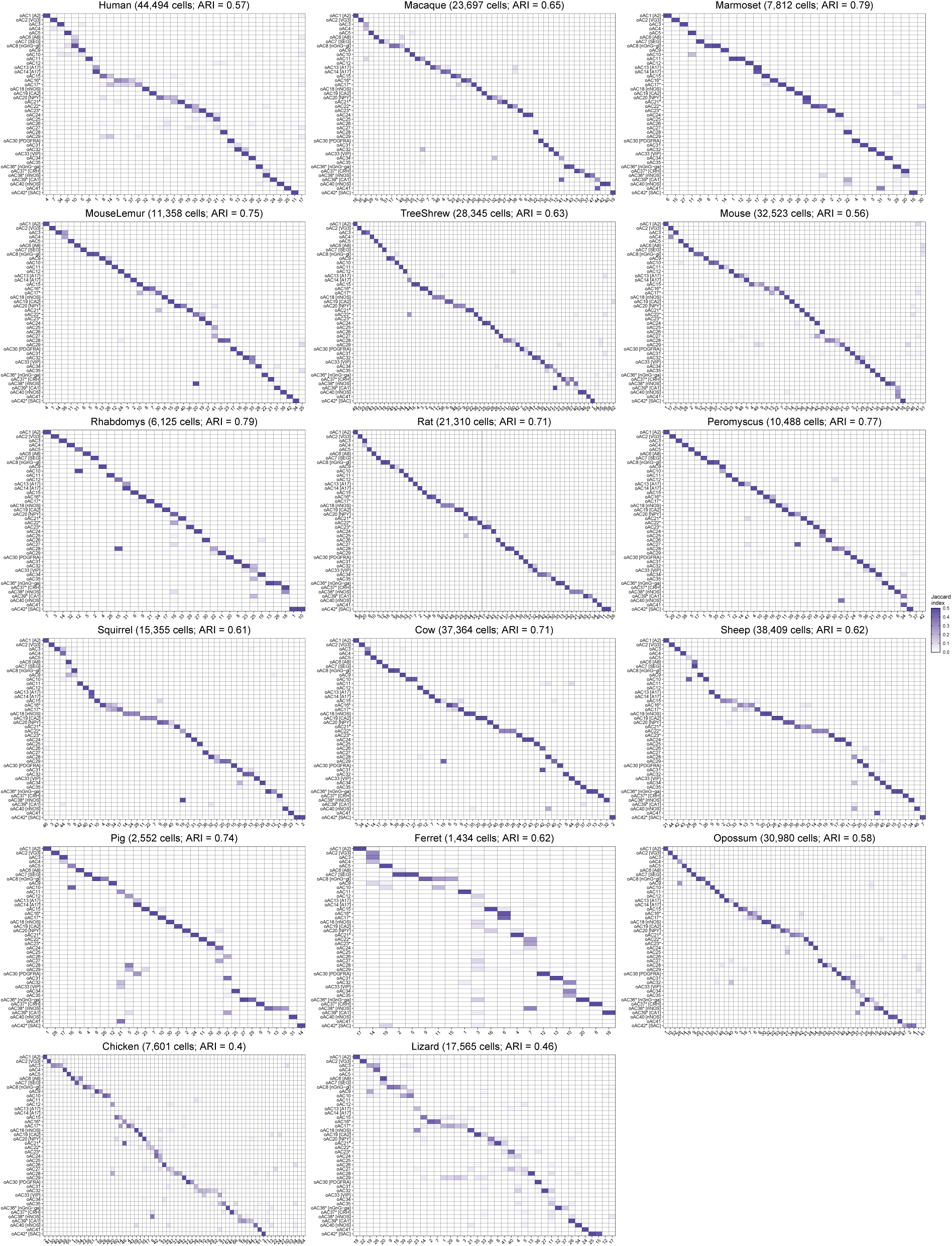
Correspondence between oACs and species-specific clusters. Heatmaps showing Jaccard overlap between oACs (y-axis) and species-specific clusters (x-axis). The diagonal structure suggests that oACs are highly conserved across amniote species. Weak correspondence in pigs and ferrets is likely due to low sample size.

**Figure S7.**
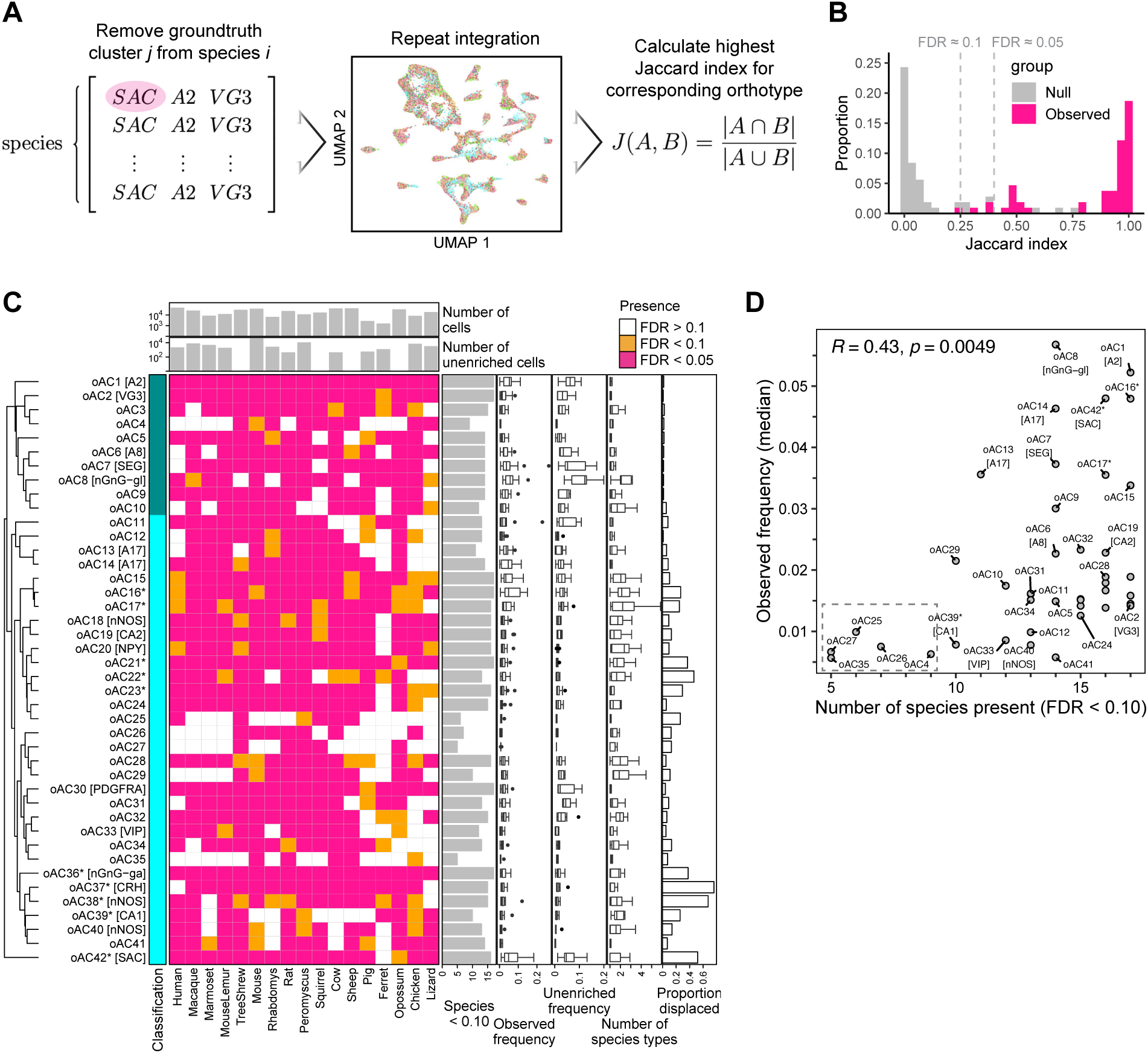
Quantifying oAC presence in species. A) Derivation of a null distribution of presence scores by iterative removal of manually annotated clusters (SAC, A2, and VG3 ACs) followed by integration and presence score computation. The presence score of an oAC in a given species is defined as the maximum Jaccard index. B) Null vs. observed distribution of presence scores. Notice that the presence scores for the null distribution are much lower, indicating that when a cell type is removed, the integration does not erroneously identify the oAC as present. The rare cases of high presence scores in the null distribution were driven by residual cells from the true cell type or recruitment of cells from a closely related cell type. C) Heatmap showing the presence of each oACs in each species. The top annotation bars shows the number of total and unenriched cells recovered for each species. The right annotation bars show metadata for the oACs. The first graph show the number of species in which the oAC was identified (p < 0.10). The next two box plots show the observed frequency (see below) and the unenriched frequency. The fourth annotation bar shows the estimated number of species types represented by the oAC, computed as the inverse Simpson index of the within-species clustering. The last bar graph shows the inferred proportion displacement for each oAC. Low frequency oACs are “absent” in several species, and poorly sampled species (e.g. ferret) exhibit several “missing” oACs. Both cases of omission likely reflect sample size limitation, and should not be interpreted as loss of cell type without further proof. D) The number of species in which an oAC is detected is significantly correlated with its observed frequency (the raw frequency observed in the atlas, which may be biased by FACS/FANS enrichment). All the oACs detected in less than 10 species at an FDR < 0.10 (oAC4, oAC25, oAC26, oAC27, oAC35; shown in grey box) are poorly sampled.

**Figure S8.**
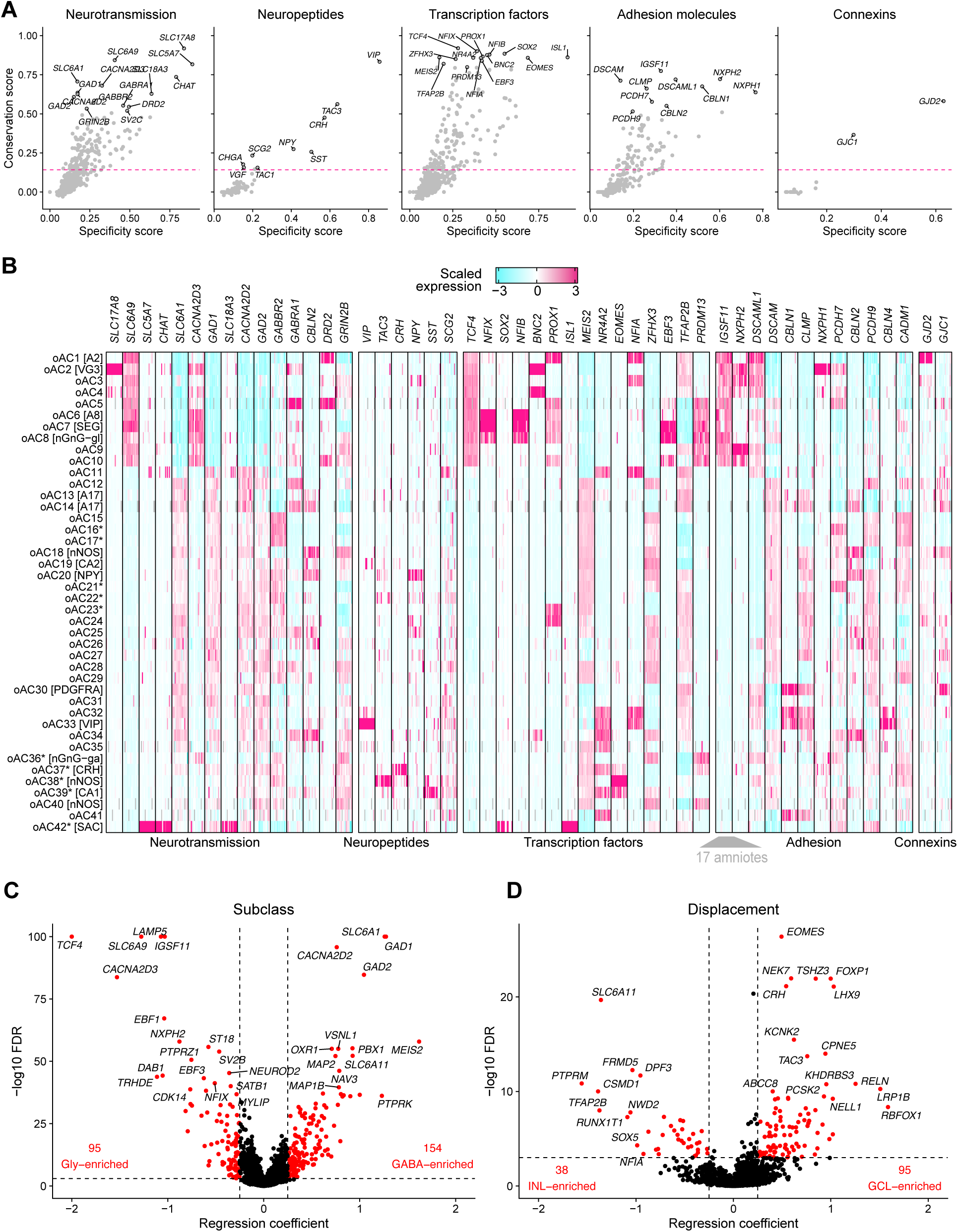
Conserved molecules involved in neurotransmission, neuropeptide signaling, transcriptional regulation, adhesion, and gap junction coupling. A) Scatterplots showing the conservation score (y-axis) for each gene versus the max specificity score (x-axis). Each dot represents one gene. See **Supplementary Materials and Methods** for descriptions of each score. Pink lines correspond to the significance threshold for the conservation score (*p* < 1.5×10^-3^ by permutation test, 17,113 permutations). B) Heatmap showing the expression of top conserved markers, stratified by species. C) Volcano plot showing genes associated with AC subclass (GABAergic versus glycinergic). Association to proportion displaced was determined by fitting a linear model for each gene, with species as a covariate. D) Volcano plot showing genes associated with proportion displaced. Association to proportion displaced was determined by fitting a linear model for each gene, with species and GABA/glycinergic subclass as a covariate.

**Figure S9.**
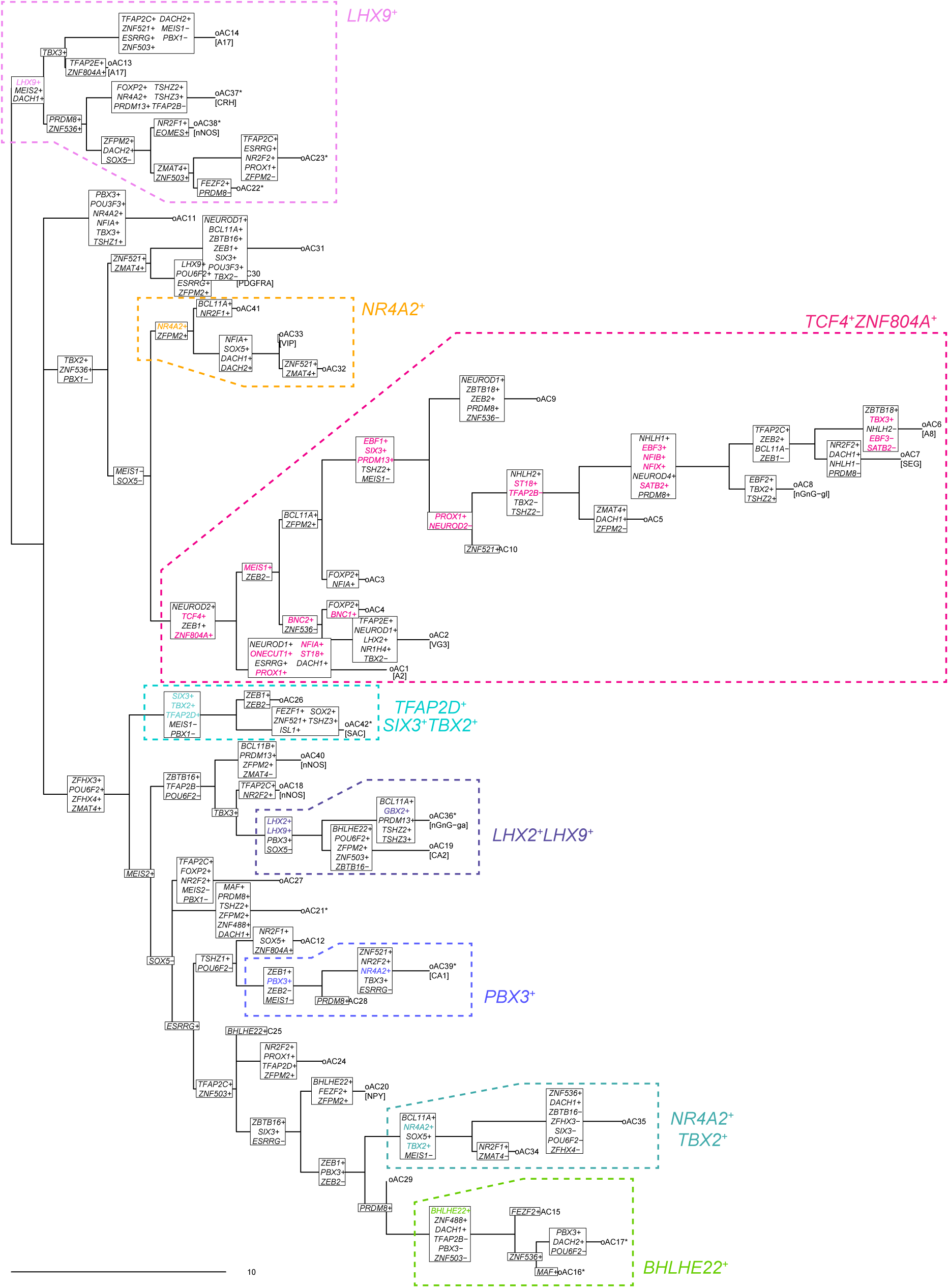
Full maximum parsimony tree with transcription factor switching events. Switching events were assigned to nodes on the tree based on the maximum parsimony reconstruction (see **Methods**). Clades shown in **Fig. 5** are highlighted by colored boxes. Genes discussed in the text are bolded and colored.

**Figure S10.**
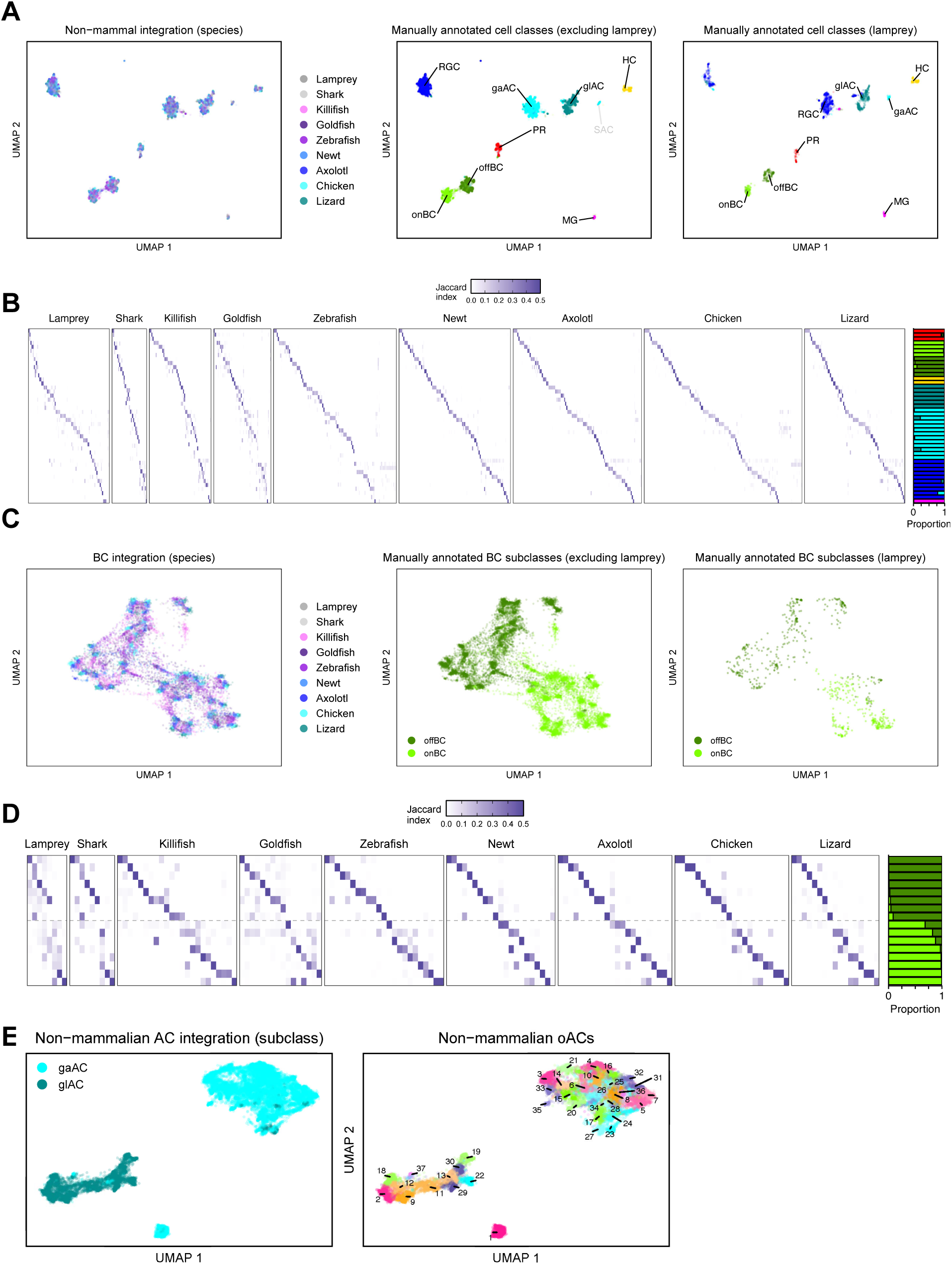
Conservation of major cell classes and bipolar cell types across 9 non-mammalian vertebrates. BCs are further split into ON and OFF types, and ACs are further split into GABAergic (gabaAC) and glycinergic (glyAC) types. A) UMAP embedding of the SAMap integration of all major cell classes (PR, BC, HC, AC, RGC, MG). Left: colored by species; middle: colored by cell class (without lamprey); right: colored by lamprey cell class. Lamprey is shown separately since many clusters annotated as RGCs by the original study cluster with ACs. B) Left: Heatmaps showing the correspondence (Jaccard index) between leiden clusters in non-mammalian integration and the within-species clustering. Right: Bar graph showing the cell class composition based on the within-species cell class annotation. C) UMAP embedding of the SAMap integration of BCs. Left: colored by species; middle: colored by cell class (without lamprey); right) colored by lamprey cell class. D) Left: Heatmaps showing the correspondence (Jaccard index) between leiden clusters from non-mammalian BC integration and within-species clustering. Right: Bar graph showing the cell class composition based on the within-species cell class annotation. E) Left: UMAP of non-mammalian AC integration, colored by GABAergic (gaAC) or glycinergic (glAC) subclass. Right: UMAP of non-mammalian AC integration, colored by non-mammalian oAC (leiden clusters).

**Figure S11.**
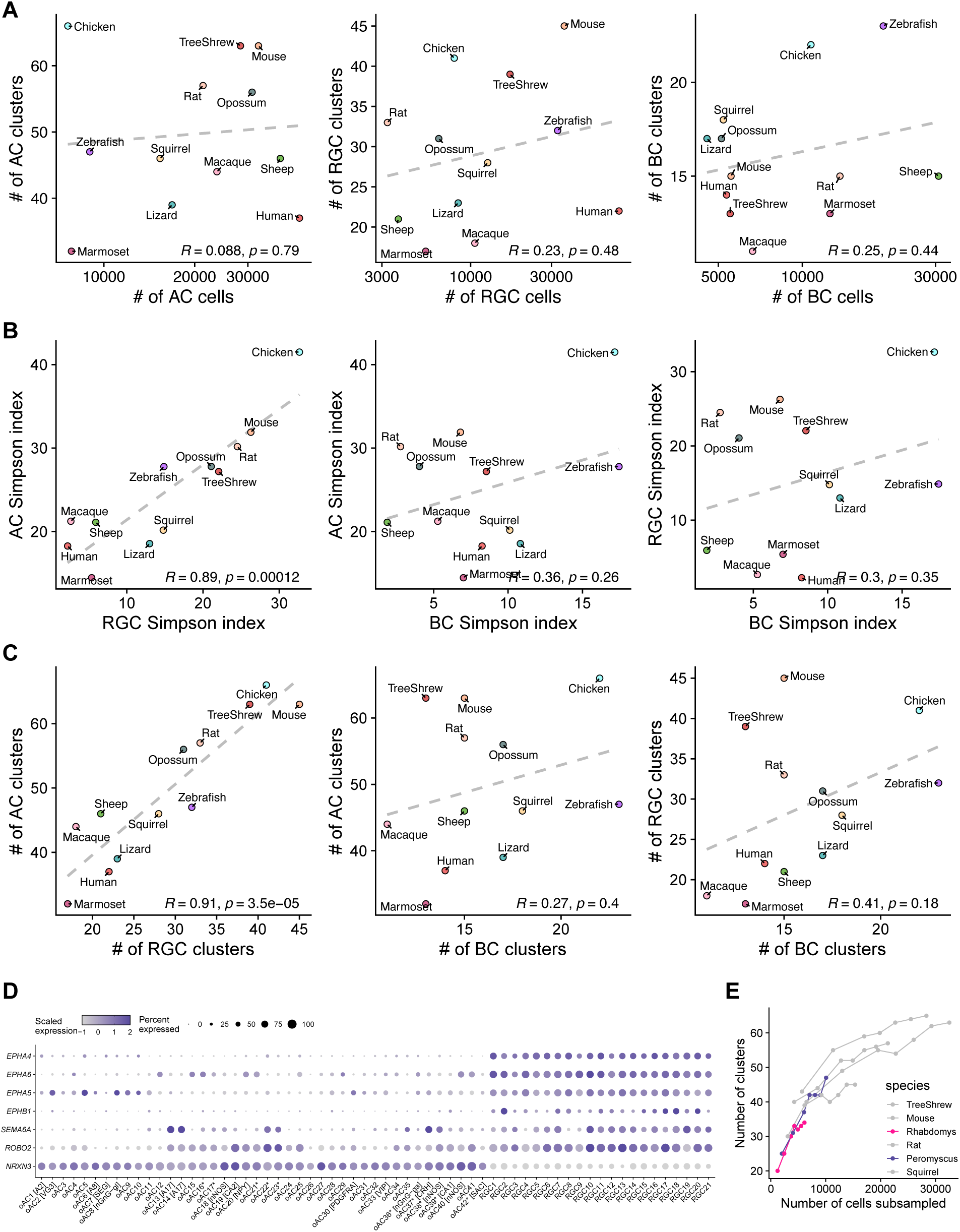
Association between AC and RGC diversity. A) No significant correlation between number of AC clusters and number of cells recovered for ACs (left), RGCs (middle), and BCs (right). Since we only used species with >3k cells of each class, there is little evidence for the presence of any sampling bias. Number of cells (x-axes) are plotted on a log-scale. B) Comparison of the diversity of AC vs. RGC, AC vs. BC, and RGC vs. BC. Diversity is quantified for each class using the Simpson diversity index (see **Methods**). Only RGC and AC diversity are significantly correlated. We only used the 12 species with greater than 3k ACs and 3k RGCs to avoid cases of undersampling. C) Comparison of the number of clusters of AC vs. RGC, AC vs. BC, and RGC vs. BCs. As in panel B, significant correlation is only observed for AC vs. RGC. D) Violin plots showing the expression of top differentially expressed adhesion molecules distinguishing oACs (this study) and oRGCs (Hahn et al., 2023). These may represent genes that are involved in directing projections to the optic nerve (RGC-enriched) or to the IPL (AC-enriched). E) Line graphs showing the number of clusters found when downsampling to 20, 40, 60, 70, 80, 90, and 100% of the cells in each rodent or scandentian dataset. Notice that rhabdomys and peromyscus have fewer total cells and the number of clusters has not reached saturation.

## Notes

### Competing Interest Statement

The authors have declared no competing interest.

